# Semiparametric Partial Common Principal Component Analysis for Covariance Matrices

**DOI:** 10.1101/808527

**Authors:** Bingkai Wang, Xi Luo, Yi Zhao, Brian Caffo

## Abstract

We consider the problem of jointly modeling multiple covariance matrices by partial common principal component analysis (PCPCA), which assumes a proportion of eigenvectors to be shared across covariance matrices and the rest to be individual-specific. This paper proposes consistent estimators of the shared eigenvectors in PCPCA as the number of matrices or the number of samples to estimate each matrix goes to infinity. We prove such asymptotic results without making any assumptions on the ranks of eigenvalues that are associated with the shared eigenvectors. When the number of samples goes to infinity, our results do not require the data to be Gaussian distributed. Furthermore, this paper introduces a sequential testing procedure to identify the number of shared eigenvectors in PCPCA. In simulation studies, our method shows higher accuracy in estimating the shared eigenvectors than competing methods. Applied to a motor-task functional magnetic resonance imaging data set, our estimator identifies meaningful brain networks that are consistent with current scientific understandings of motor networks during a motor paradigm.

## 1. Introduction

Common principal component analysis (CPCA) is an approach that simultaneously models multiple covariance matrices. It extends the idea of principal component analysis by assuming all covariance matrices share the same set of eigenvectors. Since it was first introduced by Flury (1984), CPCA has been extensively applied in various fields including statistics (Gu, 2016; Pepler et al., 2016), finance (Goyal et al., 2008; Xu et al., 2019), and computer science (Ye et al., 2012; Hadjipantelis et al., 2015).

Extensions of CPCA have been investigated from multiple angles. Flury (1987) proposed partial common principal component analysis (PCPCA), where only a proportion of the eigenvectors was assumed to be shared across covariance matrices and the rest to be individual-specific. Another direction relaxed the Gaussianity assumption in CPCA, resulting in asymptotic theory for non-Gaussian distributions (Boik, 2002; Hallin et al., 2010). Other extensions include Bayesian approaches (Hoff, 2009), algorithm acceleration (Browne and McNicholas, 2014) and modifications for high-dimensional data (Franks and Hoff, 2019). Among these extensions of CPCA, PCPCA continues to be appealing, as it relaxes the assumption of a completely common eigenspace across matrices while partially preserving the straightforward interpretation of common eigenvectors, i.e., eigenvectors shared across matrices. Related work on this topic includes Krzanowski (1984), Schott (1999), Boik (2002), Lock et al. (2013) and Pepler et al. (2016).

In spite of these extensions, some questions related to PCPCA remain unanswered. Given the number of common eigenvectors, how one can identify the common eigenvectors from a pool of eigenvectors requires further investigation. Flury (1987) assumed that “some order of the common components is defined”. Some of the literature assumed that the common eigenvectors are those associated with the largest eigenvalues across all covariance matrices (Schott, 1999; Crainiceanu et al., 2011). However, common eigenvectors may be associated with small eigenvalues, or the corresponding eigenvalue of a common eigenvector ranks differently across matrices. This question becomes more challenging if the number of common eigenvectors is unknown or the data is not Gaussian distributed. Regarding these points, Pepler et al. (2016) developed a non-parametric method to select the common eigenvectors in the special case of two covariance matrices. With multiple asymmetric matrices as the response, Lock et al. (2013) proposed a linear model to identify latent factors that explain the joint and individual data variation (JIVE) and Zhou et al. (2016) generalized JIVE to a common and individual feature extraction (CIFE) framework. None of these methods, however, studied the asymptotic properties.

In this paper, we propose a semiparametric PCPCA approach, which can consistently estimate the common eigenvectors, without making any assumptions on the ranks of eigenvalues that are associated with common eigenvectors. Our method builds on an idea from Krzanowski (1984), where a semiparametric approach was proposed in the context of CPCA. We extend this idea to semiparametric PCPCA and provide asymptotic results for our methods as the number of matrices, or the number of samples to estimate each matrix, goes to infinity (both with fixed dimension). If the number of samples goes to infinity, our results do not require the data to be Gaussian distributed. When the number of common eigenvectors is unknown, we develop a sequential testing procedure, which effectively controls the type I error for Gaussian distributed data. As shown in the simulation study, our method outperforms existing methods in estimating the common eigenvectors in a variety of scenarios.

In the next section, we introduce PCPCA. In Section 3, we present our proposed semi-parametric method to identify the common eigenvectors. We evaluate the performance of our proposed method through simulation studies in Section 4. An application to an fMRI data set is provided in Section 5. Section 6 summarises this paper and discusses future directions.

## 2. Model and assumptions

We consider a data set, {**y**_*it*_}, for *t* ∈ {1, …, *T*} and *i* ∈ {1, …, *n*}, where **y**_*it*_ ∈ ℝ^*p*^ are independent and identically distributed random samples from a *p*-dimensional distribution with mean zero and covariance matrix **Σ**_*i*_. In our application example, **y**_*it*_ is a sample of brain fMRI measurements of *p* regions from subject *i* at time point *t*. We assume that **Σ**_*i*_ satisfies the following partial common principal component (PCPC) model:

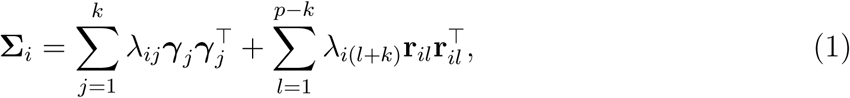

where 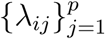 are the eigenvalues of covariance matrix **Σ**_*i*_ and *k* is the largest integer such that formulation 1 holds. The ***γ***_*j*_, for *j* = 1, …, *k*, are the unit-length common eigenvectors across subjects. Let **Γ** = (***γ***_1_, …, ***γ***_*k*_) ∈ ℝ^*p×k*^ (*k* ≤ *p*) be the orthonormal matrix of the common eigenvectors. The **r**_*il*_, for *l* = 1, …, *p* − *k*, are unit-length individual-specific eigenvectors of subject *i*. Let **R**_*i*_ = (**r**_*i*1_, …, **r**_*i,p*−*k*_) ∈ ℝ^p×(*p*−*k*)^ be the orthonormal matrix of the individual-specific eigenvectors. We assume that **R**_*i*_ is orthogonal to **Γ**, i.e., **Γ**^T^**R**_*i*_ = **0**. Let **Λ**_*i*_ = diag{*λ*_*i*1_, …, *λ*_*ik*_} and 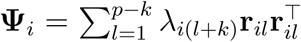. Then, the PCPC model (1) can be reformulated as:

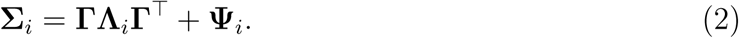

First proposed by Flury (1987), the PCPC model has an interpretation analogous to CPCA. A CPC, defined by 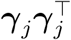 for *j* = 1, …, *k*, is shared across all matrices. We emphasize that our definition of CPC is different from Flury (1984) (or the principal component in PCA) where a CPC is defined as 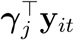, since we focus on the shared covariance structure across matrices instead of individual-specific eigenvalues. For example, in our application, a CPC represents a functional brain network in the sense that it represents correlations in functional brain measures consistent across subjects. The corresponding diagonal entry of **Λ**_*i*_ is interpreted as the variation of the CPC in subject *i*. On the other hand, **Ψ**_*i*_ is the individual-specific model component, which varies across subjects. A toy example of the PCPC model with *p* = 4 and *k* = 2 is shown in Figure 1.

**Figure 1.**
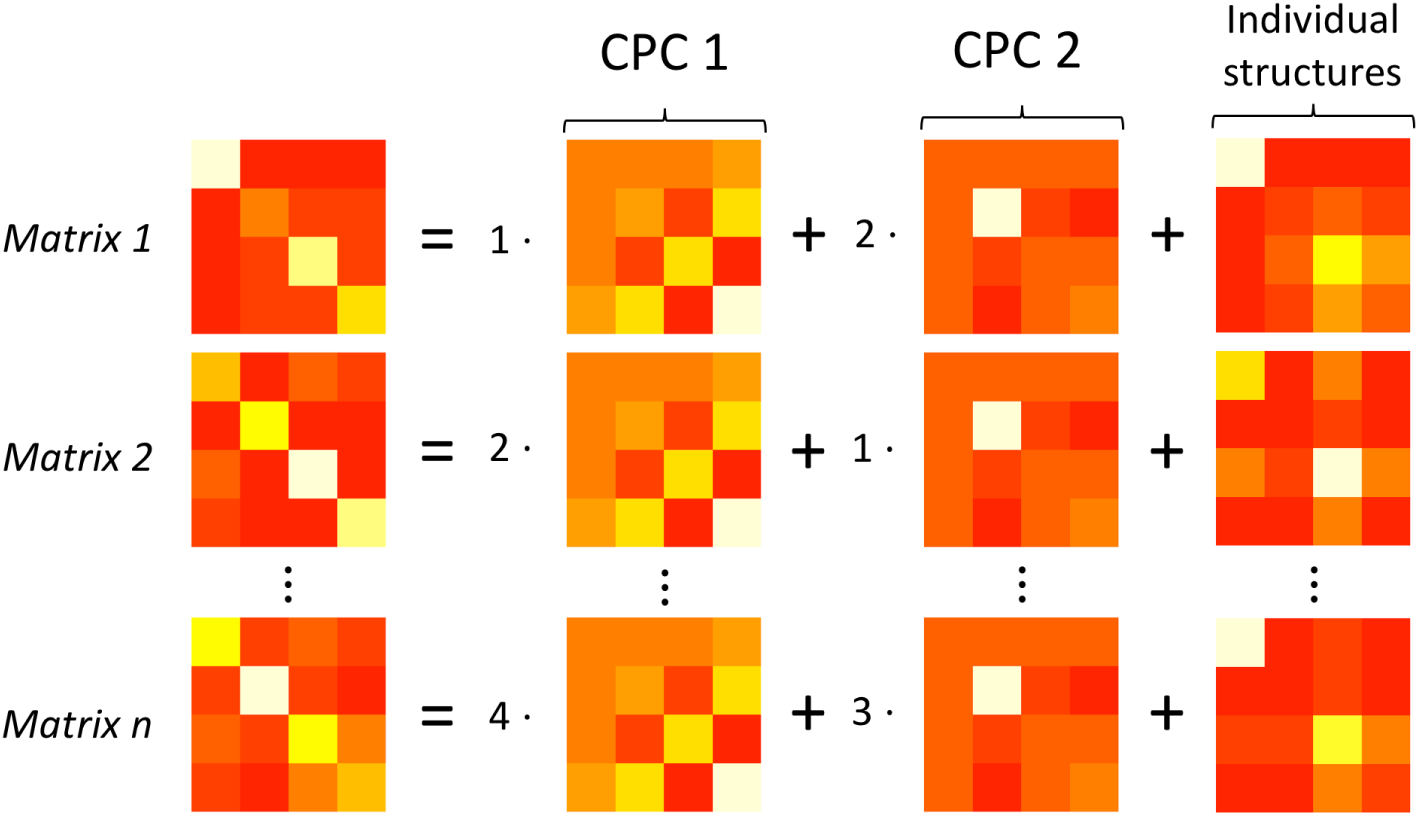
An example of the PCPC model. Each covariance matrix consists of two CPCs and an individual structure. Each CPC has rank 1 and norm 1.

Our goal is to both find *k* and estimate **Γ** consistently, as either *n* → ∞ or *T* → ∞, with *p* fixed. When estimating **Γ**, existing methods, such as Schott (1999) and Crainiceanu et al. (2011), assumed that CPCs are associated with the largest eigenvalues across all covariance matrices. This is a restrictive assumption, since the corresponding eigenvalue of a CPC may rank consistently low or differently across matrices. For instance, our toy example in Figure 1 shows that CPCs are associated with small eigenvalues in matrices 1 and 2, but with large eigenvalues in matrix *n*. In addition, in many scientific applications, there is no priori reason to assume that the variation explained by the common components dominates the variation explained by the individual-specific components. For this reason, we do not make any assumptions regarding the rank of CPC-related eigenvalues.

The PCPC model shares some common features with existing partial information decomposition methods, but there exist major differences. Crainiceanu et al. (2011) provided a population value decomposition (PVD) model where common eigenvectors are extracted from concatenated individual eigenvectors. This procedure presumes that common inter-subject components are associated with the largest individual eigenvalues. Moreover, the approach does not consider group level diagonalization as a goal. Lock et al. (2013) introduced the JIVE model, which decomposes information from multiple data sources into common components and individual components. Zhou et al. (2016) generalized the JIVE model by a CIFE framework, which has the same objective function as JIVE. Unlike the PCPC model, the common components identified by JIVE and CIFE are not unique, which can make them hard to interpret in practice. Furthermore, PVD, JIVE and CIFE are empirical methods with no asymptotic guarantees.

More recently, Wang et al. (2019) proposed a common reducing subspace model, which assumes 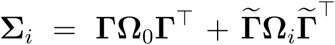, where 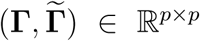 forms an eigenbasis and **Ω**_0_ ∈ ℝ^*k×k*^, **Ω**_*i*_ ∈ ℝ^(*p*−*k*)*×*(*p*−*k*)^, *i* = 1, …, *n* are positive definite matrices. This model can be reformulated as **Σ**_*i*_ = **ΓΛΓ**^T^ + **Ψ**_*i*_, where **Λ** ∈ ℝ^*k×k*^ is a positive definite diagonal matrix shared across *i* and **Ψ**_*i*_ ∈ℝ^*p×p*^ is a positive semi-definite matrix orthogonal to **Γ** with rank *p* − *k*. Compared with the PCPC model (1), this model requires **Λ**_*i*_ *≡* **Λ**, and is hence a special case of the PCPC model.

To achieve the identifiability of **Γ** and the consistency of our proposed estimator, for the PCPC model (1), we impose the following assumptions for the asymptotics when *n* → ∞.

**Assumption A (for *n* → ∞):**

1. *T* and *p* are fixed with *T >* 0 and *p >* 1.
2. Each (*λ*_*i*1_, …, *λ*_*ik*_), *i* ∈ {1, …, *n*}, is an independent and identically distributed random sample from a distribution with finite mean 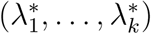 and finite variance. Furthermore, elements of (*λ*_*i*1_, …, *λ*_*ik*_) are independent of each other.
3. Each **Ψ**_*i*_, *i* ∈ {1, …, *n*}, is an independent and identically distributed random sample from a distribution with finite mean **Ψ**^*\**^ and finite second-order moment. Both **Ψ**_*i*_ and **Ψ**^*\**^ are symmetric positive semi-definite matrices with rank *p* − *k* and are orthogonal to **Γ**.
4. The matrix **ΓΛ**^*\**^**Γ**^T^ + **Ψ**^*\**^ has distinct eigenvalues, where 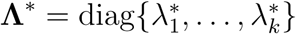.
5. For each *i* ∈ {1, …, *n*}, **y**_*it*_ is normally distributed given **Σ**_*i*_.

To the best of our knowledge, we are the first to provide asymptotic results as the number of matrices goes to infinity. Different from the literature where *n* is fixed (Flury, 1987; Boik, 2002; Pepler et al., 2016; Wang et al., 2019), Assumptions A (2) and (3) assume (*λ*_*i*1_, …, *λ*_*ik*_) and **Ψ**_*i*_ are random variables instead of fixed parameters, since otherwise, the number of parameters would explode as *n* increases. Assumption A (4) is required for identifiability of **Γ** in matrix perturbation theory. The Gaussian assumption in Assumption A (5) is made for convenience and is stronger than required for our results. For the proof, we only need the fourth-order moment of **y**_*it*_ to be the same as the fourth-order moment of a Gaussian distribution, with mean 0 and covariance **Σ**_*i*_.

In some cases, *n* is small but *T* is large. For example, in fMRI data analysis, the number of subjects may be small, but subjects may have long fMRI scans. In other measures with rapid sampling, such as electroencephalograms, this is frequently the case. For such data sets, we prove a similar asymptotic theory as *T* → ∞ with *n* and *p* fixed. This asymptotic theory requires the following assumptions.

**Assumption B (for *T* → ∞):**

1. *n* and *p* are fixed with *n >* 1 and *p >* 1.
2. For each *i* ∈ {1, …, *n*}, (*λ*_*i*1_, …, *λ*_*ik*_) and **Ψ**_*i*_ are fixed.
3. The eigenvalues of 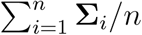 are distinct.
4. The fourth-order moment of **y**_*it*_ is bounded for *i* = 1, …, *n*.

Assumption B (2) implies that the asymptotics are conditional on **Σ**_*i*_, *i* = 1, …, *n*. The reason to pursue conditional asymptotics is that *n* is fixed and inference on these specific *n* distributions is of interest. Unlike Assumption A where a Gaussian distribution is assumed, Assumption B is semiparametric, since it does not put constraints on higher-order moments, except that the fourth-order moment is bounded. Compared with existing asymptotic results for *T* → ∞, Assumption B is weaker. Flury (1987) and Schott (1999) both assumed that {**y**_*it*_} are normally distributed. Boik (2002) provided asymptotic results for non-normal data, but modeled eigenvalues as known smooth functions of parameters. Though Pepler et al. (2016) and Hallin et al. (2010) relaxed Assumptions B (3) and (4), the former work only focused on the case when *n* = 2 and the latter was for CPCA.

In addition to Assumption A or Assumption B, we also assume that {**y**_*it*_}, *t* = 1, …, *T* are independent of each other. However, in many real-world applications, such as our fMRI data example, {**y**_*it*_} can be temporarily correlated. We consider a generic constraint on the temporal correlation that, for 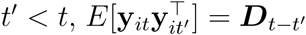 where ***D***_*t*−*t*_′ is a diagonal matrix and ***D***_*t*−*t*_′ = **0** if *t > t*′ + *c* for some constant *c*. This assumed constraint is satisfied for many time series models, including Bickel and Gel (2011) and Guo et al. (2016), and approximately satisfied under the auto-regressive model. Under this assumption, our theoretical results for *n* → ∞ still hold. Alternatively, when *T* → ∞, under the same assumption, one can adopt an auto-regressive moving-average (ARMA) model for pre-whitening {**y**_*it*_} to remove temporal dependence, which is commonly used in fMRI data analysis (Lindquist et al., 2008; Olszowy et al., 2019).

## 3. Estimation

In this section, we introduce our estimation procedure under two scenarios: (1) the number of CPCs is known and (2) the number of CPCs is unknown. When the number of CPCs is known, we prove that our proposed estimator of the common eigenvectors is consistent. When the number of CPCs is unknown, the estimation procedure has two steps: we first use a sequential testing procedure to estimate the number of CPCs, and then calculate our proposed estimator using the estimated number of CPCs.

### 3.1 The number of CPCs is known

When the number of CPCs is known, we propose to estimate **Γ** in two steps: first getting *p* CPC candidates for **Γ**, denoted as 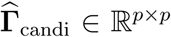, and then selecting *k* columns from 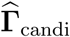 as 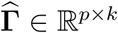.

In the first step, 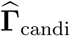 is calculated as the eigenvectors of 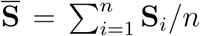, where 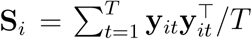 is the sample covariance matrix of subject *i*. The columns of 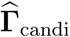 are ordered in a way that the corresponding eigenvalue of each eigenvector is decreasing. We first define the consistency of an eigenvector estimator.

Definition 1: Let {**x**_*s*_ : *s* = 1, 2, …} denote a series of random vectors in ℝ^*p*^ with 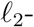 norm 1; that is ‖**x**_*s*_ ‖_2_ = 1 for all *s*. Let **x** be a vector in ℝ^*p*^ such that ‖**x**‖_2_ = 1. As *s* → ∞, **x**_*s*_ is consistent to **x** if

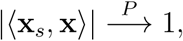

where ⟨·, ·⟩ is the inner product defined in ℝ^*p*^ and 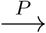 denotes convergence in probability.

Under Definition 1, the following theorem shows that *k* out of *p* columns of 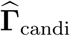 are consistent estimators of the columns of **Γ** as *n* or *T* goes to infinity, which is a direct generalization of spectral properties of 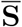.

Theorem 1: *Assume the PCPC model* (1) *holds.*

1. *Under Assumption A, for any column* ***γ***_*j*_ *of* **Γ** *(j* = 1, …, *k), there exists a column of* 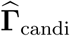 *that is consistent to* ***γ***_*j*_ *as n* → ∞. *Explicitly, let* **e**_*l*_ ∈ℝ^*p*^ *denote a p-dimensional vector with the l-th entry one and rest zero, then, there exists l*(*j*) ∈ {1, …, *p*}, *such that*

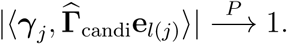
2. *Under Assumption B, for any column* ***γ***_*j*_ *of* **Γ** *(j* = 1, …, *k), there exists a column of* 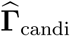 *that is consistent to* ***γ***_*j*_ as *T* → ∞.

Theorem 1 implies that, by properly ordering the columns of 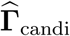, we can achieve that the *j*-th column of 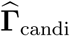 converges in probability to ***γ***_*j*_ for *j* = 1, …, *k*. To find this ordering, we define a deviation from commonality metric for each column of 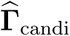:

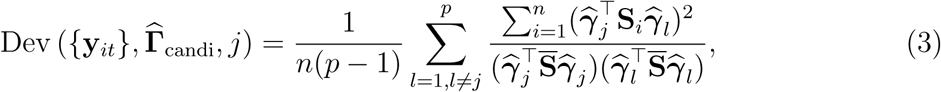

where 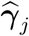 is the *j*-th column of 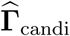.

For the deviation from commonality metric, we expect it to be small if 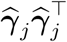 is close to a true CPC and large otherwise. For illustration, we assume *T* is large enough such that the sample estimates can be replaced by their population targets; that is, 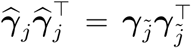 for some 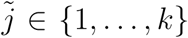 and **S**_*i*_ = **Σ**_*i*_. Then the PCPC model (1) implies that 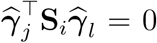 for *l ≠ j* and hence Dev 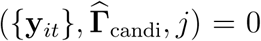. If 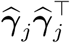 and 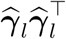 are not close to any CPC, then 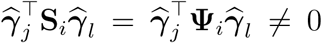 for some *i* and hence Dev 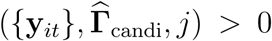. In general, on the right-hand side of Equation (3), the numerator captures the sum of the squared (*j, l*) element in 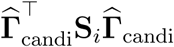 and the denominator is a normalizing term that eliminates the effect of magnitude difference in the eigenvalues. The following theorem shows that Dev 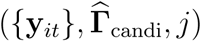 can be used to order the columns of 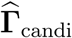 and estimate **Γ**.

Theorem 2: *For all j* ∈ {1, …, *k*}, *let* 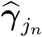 *be the estimate of* ***γ***_*j*_ *in* 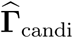, *where* 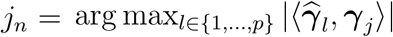. *Assume the PCPC model* (1) *holds.*

1. *Under Assumption A, as n → ∞, we have*

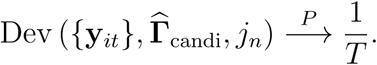

In addition, let L_*n*_ = {1, …, *p*} \ {1_*n*_, …, *k*_*n*_}, *then there exists a positive constant C independent of n, such that, as n* → ∞,

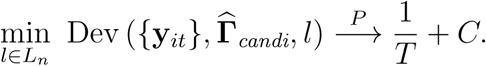
2. *Under Assumption B, as T* → ∞, we have

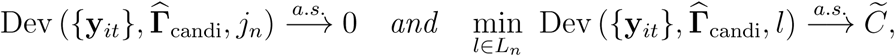

*where 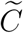 surely. is a positive constant independent of T and 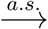 denotes convergence almost surely.*

Given Theorem 2, in practice, we can rank the columns of 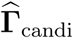 in increasing order of 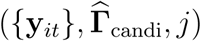 for *j* = 1, …, *p* and select the first *k* columns as 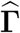. If 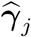 is selected, we call 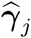 a common eigenvector estimate and 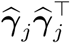 a CPC estimate.

In Theorem 2, the asymptotic results of *n* → ∞ and *T* → ∞ are different, which results from the different assumptions made in the two cases. When *n* → ∞, the deviation from commonality metric is related to the fourth-order moment of **y**_*it*_, which yields a positive probability limit for a CPC estimate. When *T* → ∞, we have 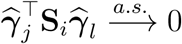 for *l ≠ j* if and only if 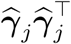 is a CPC estimate, making the deviation from commonality metric converge to 0 only for a CPC estimate. Despite these differences, CPC estimates in both cases have the least deviation from commonality metric among all columns of 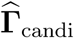 asymptotically, which is essential for identifying CPC estimates from 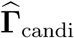.

When *n* and *T* are small, some columns of 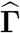 may have a large deviation from commonality metric and are not “close” to any CPC. This bias, however, will disappear as *n* → ∞ or *T* → ∞, as guaranteed by Theorems 1 and 2. While our theorems hold for all *k* ∈ {0, 1, …, *p* − 2, *p*}, the convergence rate can be faster for larger *k*. Under Assumption A, when *k* is large, {**y**_*it*_} for different *i* share more in common, which reduces the variability of the eigenvectors of 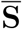. Under Assumption B, as *k* increases, the number of parameters in the PCPC model (1) decreases, and the effective sample size to estimate each parameter increases. We leave the study of the convergence rate as a function of *p* and *k* to future research. The estimating procedure is summarized in Algorithm 1.

#### Algorithm 1 An algorithm to estimate CPCs in model (1) when *k* is known

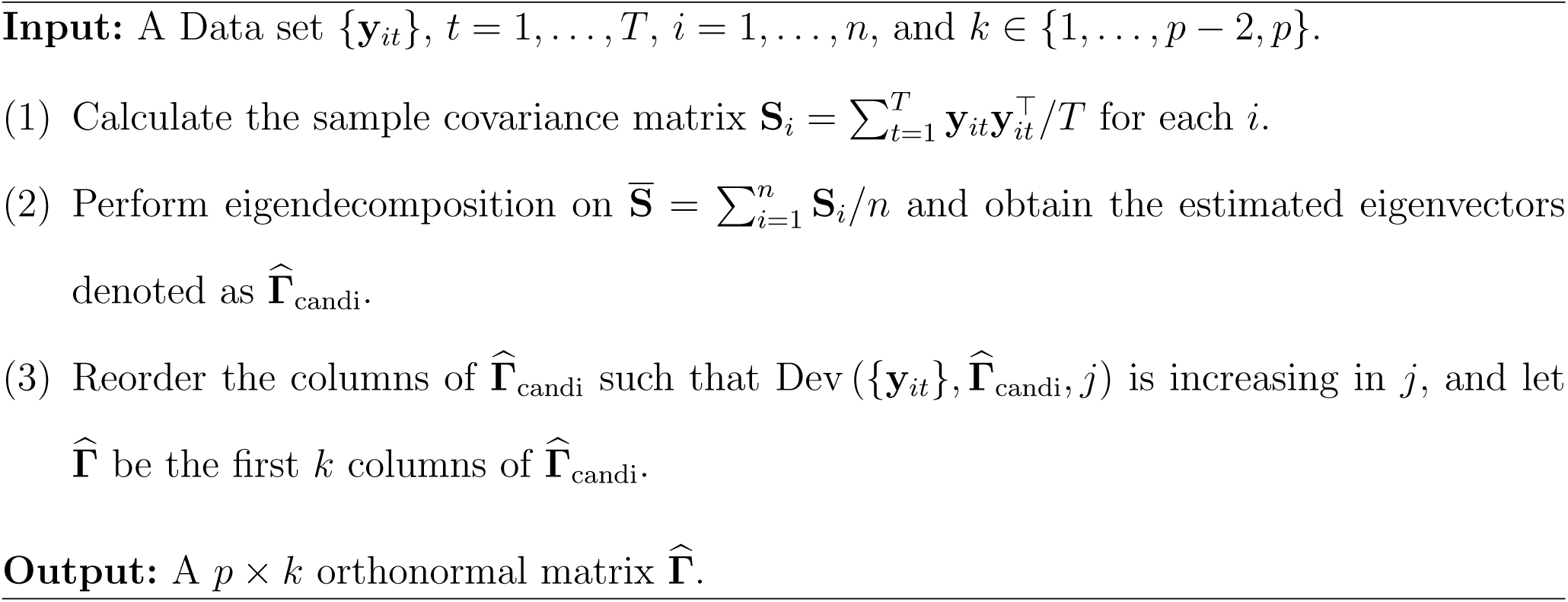

With the number of CPCs given, we generalize the results of Flury (1987) in three directions. First, Algorithm 1 can consistently estimate CPCs as *n* → ∞, a case Flury (1987) did not cover. Second, when *T* → ∞, Theorems 1 and 2 relax the Gaussian assumption made by Flury (1987). Third, Theorems 1 and 2 guarantee the identification of the CPCs without making assumptions on the ranks of CPC-related eigenvalues.

Theorems 1 and 2 allow that *p > T* when *n* → ∞. When implementing Algorithm 1, the only condition is that 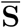 is positive definite, which is generally true if *p < nT*. However, a large *p* may substantially increase the computational complexity and affect finite-sample accuracy, as discussed in Sections 3.3 and 4, respectively.

### 3.2 The number of CPCs is unknown

Based on the idea of Schott (1999), we use a sequential hypothesis testing approach to find *k*. For *j* = 0, 1, …, *p* − 2, we sequentially perform the following testings

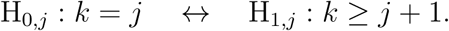

Starting from *j* = 0, if H_0,*j*_ is rejected, then we proceed to test H_0,*j*+1_; otherwise we estimate 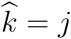. Before the first test, we order the columns of 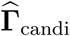 such that Dev 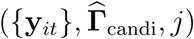 is increasing in *j*. When testing the *j*-th hypothesis, we simulate the distribution of Dev 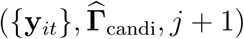 under H_0,*j*_, denoted as 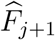, and reject H_0,*j*_ if Dev 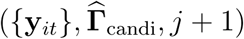 is smaller than the *α*-quantile of 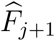. The logic of this rejection rule is that Dev 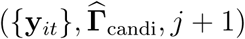 is small under H_1,*j*_, but the *α*-quantile of 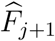 is generally large, since 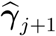 is not a common eigenvector under the null hypothesis. Adjusting for multiple testing is unnecessary here, since the family-wise type I error is 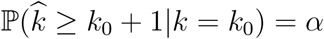, if the truth is *k* = *k*_0_.

Given **Γ** and 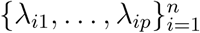 defined in the PCPC model (1), we calculate 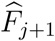 by repeating the following steps for *m* times. In practice, we can approximate **Γ** using 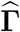 output in Algorithm 1, estimate {*λ*_*i*1_, …, *λ*_*ij*_} by diagonal entries of 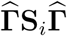 and estimate {*λ*_*i*(*j*+1)_, …, *λ*_*ip*_} by the non-zero eigenvalues of 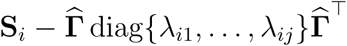. We emphasize that, different from Algorithm 1, where *p* can be larger than *T*, the above approximations are valid when *p* ≤ *T*.

1. For each *i* = 1, …, *n*, independently and uniformly generate 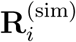 from the sample space 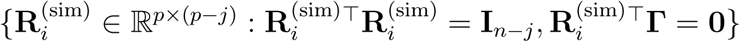.
2. Construct 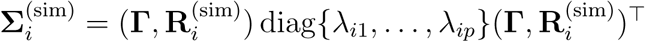. Generate 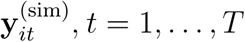, from multivariate Gaussian distribution with mean **0** and covariance 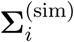.
3. Given the data set 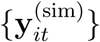, calculate 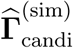 as described in Algorithm 1 and output Dev 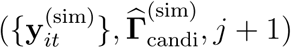.

Then 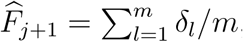, where *δ*_*l*_ denotes a point mass at Dev 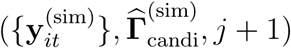 output by the *l*-th simulation. The following theorem shows that the type I error rate for each test is bounded by *α* under regularity assumptions.

Theorem 3: *Assume the PCPC model (1) holds*, {**y**_*it*_|**Σ**_*i*_} *follows a multivariate Gaussian distribution with mean* **0** *and covariance* **Σ**_*i*_, and **R**_*i*_ follows a uniform distribution on its sample space defined in Section 2. Then under H_0,*j*_, *as m* 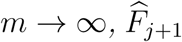 *converges in distribution to the true distribution of* Dev 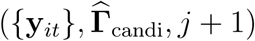 *given* **Γ** *and* {*λ*_*i*1_, …, *λ*_*ip*_}, *i* = 1, …, *n*.

A key assumption in Theorem 3 is that data are Gaussian distributed. When this assumption does not hold, the sequential testing procedure tends to be conservative, i.e. 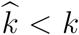, since the 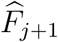 is likely to underestimate the mean and deviation of the distribution of Dev 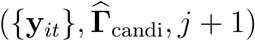. In practice, an ad hoc solution is to let 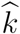 be the smallest *j* such that 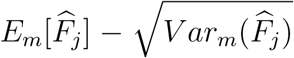 is smaller than Dev 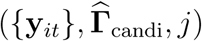, where *E*_*m*_ and *V ar*_*m*_ represent sample average and variance respectively. This solution shares the central idea of gap statistics in Tibshirani et al. (2001), which is used to determine the number of clusters in clustering. The procedure of finding *k* and estimating **Γ** is described in Algorithm 2.

Another method to find *k* is to use the hierarchy of partial chi-squared statistics proposed by Flury (1987, 1988). A nice summary of these statistics can be found in Pepler et al. (2016). The relevant application of this hierarchy in PCPCA is testing *k* = *k*_1_ *↔ k* = *k*_2_. However, this approach has two limitations. First, a set of common eigenvector estimates must be prespecified to implement the test, which is unknown under our setting since CPCs can rank differently among matrices. Second, the chi-squared test is valid only as *T* → ∞, which is a case where our approach also applies. Hence, we do not consider this method to find *k* in the simulation studies and data application.

#### Algorithm 2 A two-step algorithm to estimate CPCs in model (1) when *k* is unknown

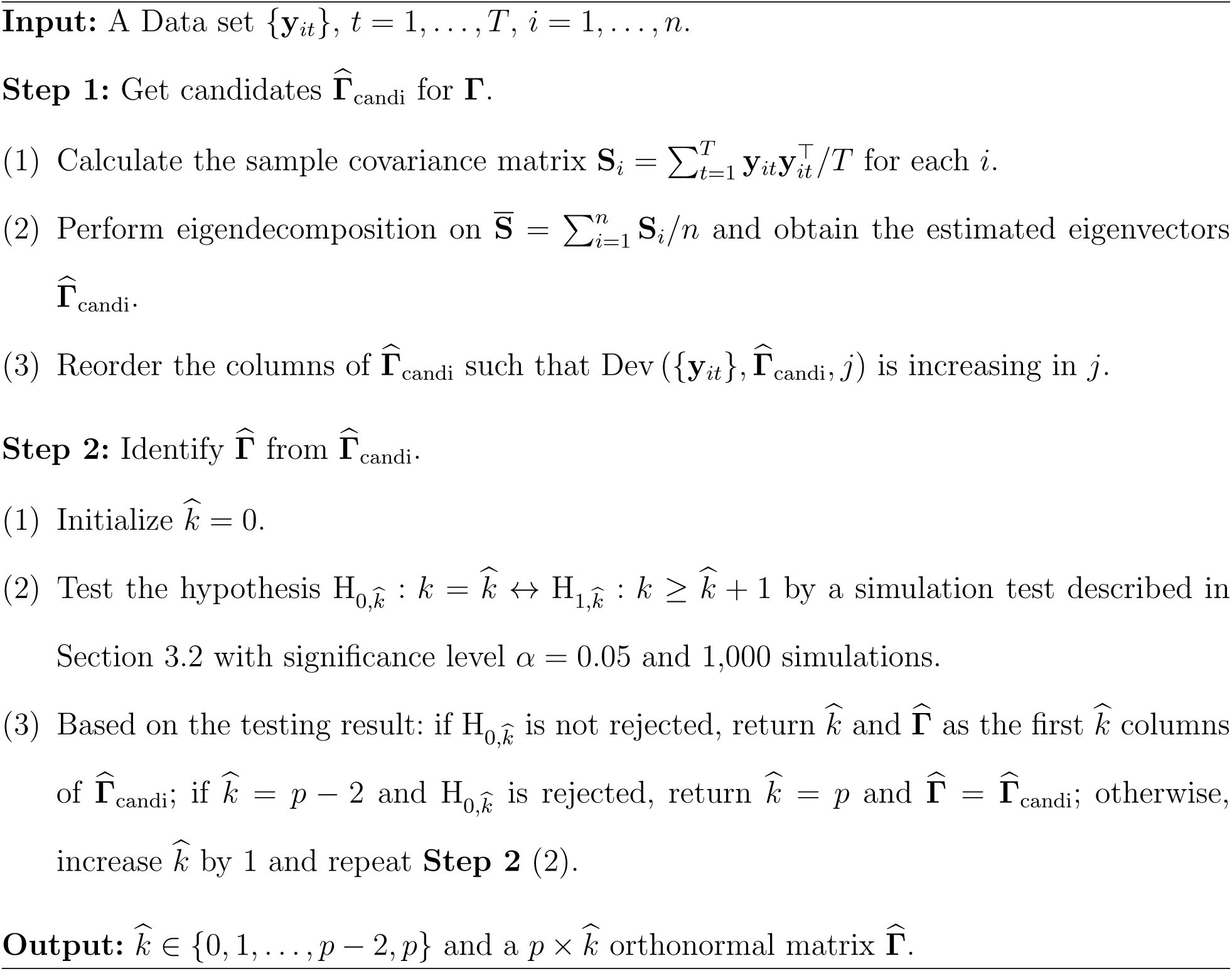

### 3.3 Computational complexity

Given parameters *k, m, n, T* and *p*, the computational complexity is *O*{*np*^2^(*p* + *T*)} for Algorithm 1 and 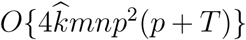 for Algorithm 2, where 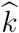is the estimate of by *k* Algorithm 2. It is straightforward to see that the dimension of matrices *p* drives the computational complexity at a rate of *p*^3^, if *T* is not too large. Furthermore, finding *k* can dramatically increase the run time if *k* and *m* are large.

As a benchmark for actual run time, we set *k* = *p* = 20, *m* = *n* = *T* = 100 and ran both algorithms on an Intel I5-8259U 2.3GHz processor in R software for 10 times. On average, Algorithm 1 took 0.04 seconds and Algorithm 2 took 453.01 seconds, where the difference is the run time due to the iterations for estimating *k*. In comparison, Flury’s algorithm (Flury and Gautschi, 1986) for CPCA, which assumes *k* is known, took 5.23 seconds under the same setting, which is roughly 100 times slower than Algorithm 1. In practice, one could reduce the run time of Algorithm 2 by parallel programming and improving code efficiency.

## 4. Simulation study

In this section, we perform three simulation studies. The first confirms the asymptotic results given by Theorem 2. The second tests the performance of Algorithms 1 and 2 under various settings. The last compares our proposed method with existing approaches under different scenarios.

### 4.1 Design and data generating mechanism

In the first simulation, we let *p* = 20 and *k* = 10. Define *λ*_*j*_ = *e*^0.5(*p*−*j*)^ for *j* = 1, …, *p*, and assume that {**y**_*it*_} follows a multivariate Gaussian distribution and CPCs rank randomly in each covariance matrix. For the study of the asymptotics as *n* → ∞, we set *T* = 50 and *n* = 50, 100, 500, 1000; and for the study of the asymptotics as *T* → ∞, we set *n* = 50 and *T* = 50, 100, 500, 1000. For each combination of *n* and *T*, we simulate data and compare the distribution of the *k*-th smallest deviation from commonality metric, which is the largest metric of CPC estimates, with the distribution of the (*k* + 1)-th smallest deviation from commonality metric, which is the smallest metric of non-CPC estimates.

The second simulation is the same as the first one, except that we consider combinations of different settings: (1) *n* = *T* = 15, 30, 100, (2) *k* = 1, 10, 20 and (3) {**y**_*it*_} follows a multivariate Gaussian distribution versus Gamma distribution. For each combination, we simulate data for 1000 times, run Algorithm 1 to get 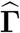, and run Algorithm 2 to get 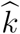 for each simulated data set. To measure the performance of Algorithm 1, we define 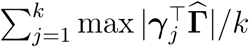 as the accuracy metric of 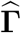. This metric lies in [0, 1] with larger values indicating better accuracy. To evaluate the sequential testing procedure, we report 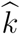 and compare it with the true *k*.

The last simulation compares our proposed method (with or without *k* known, i.e., Algorithm 1 or Algorithm 2) with Flury’s method (Flury, 1987) and the PVD method (Crainiceanu et al., 2011) through 4 scenarios below. By “Flury’s method”, we mean first running the algorithm given by Flury and Gautschi (1986) to estimate 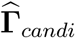 in CPCA and then selecting *k* columns associated with the largest eigenvalues of 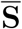. Although Flury (1987, 1988) proposed a method to estimate *k* in PCPCA, we do not implement it here, because the order of CPCs is unknown, as discussed in Section 3.2. For the PVD, we use the default setting; that is, first calculating the top *k* eigenvectors of **S**_*i*_ (denoted as **U**_*i*_) and then estimating **Γ** as the top *k* eigenvectors of **U** = (**U**_1_, …, **U**_*n*_). There are other partial information decomposition methods, such as JIVE and CIFE described in Section 2, but they do not have unique CPC estimates, which makes the comparison with these methods via simulation infeasible.

Scenario 1: {**y**_*it*_} follows a Gaussian distribution with large *n* and *T*. CPCs are associated with the largest eigenvalues in each covariance matrix.

Scenario 2: {**y**_*it*_} follows a Gaussian distribution with large *n* and *T*. The CPC-associated eigenvalues rank randomly in each covariance matrix.

Scenario 3: {**y**_*it*_} follows a Gamma distribution with large *n* and *T*. CPCs are associated with the largest eigenvalues in each covariance matrix.

Scenario 4: {**y**_*it*_} follows a Gaussian distribution with small *n* and *T*. CPCs are associated with the largest eigenvalues in each covariance matrix.

Scenario 1 serves as the reference case, where the underlying assumptions of all 4 methods are satisfied. Different from Scenario 1, Scenario 2 has randomly ranked CPC-associated eigenvalues, Scenario 3 has Gamma data generating distribution and Scenario 4 has small sample size. For each of the 4 scenarios, we consider two cases: *p* = 20 with *k* = 10 and *p* = 100 with *k* = 20, which represent small-scale and large-scale problem, respectively. When *p* = 20, we set *n* = *T* = 100 for Scenarios 1-3 and *n* = *T* = 30 for Scenario 4 and define *λ*_*j*_ = *e*^0.5(*p*−*j*)^ for *j* = 1, …, *p*. When *p* = 100, we set *n* = *T* = 1000 for Scenarios 1-3 and *n* = *T* = 150 for Scenario 4 and define *λ*_*j*_ = *e*^0.1(*p*−*j*)^ for *j* = 1, …, *p*. Similar to simulation 2, we use 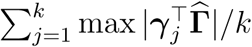 as the accuracy metric of 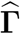.

For all simulations, if {**y**_*it*_} follows a Gaussian distribution and CPC-associated eigenvalues rank randomly, we simulate the data as follows for 1000 replications. Given *p, n, T, k* and 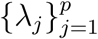, we sample one **Γ** from the space {**Γ : Γ^T^Γ = I_*k*_**} as the common eigenvectors, and randomly partition 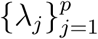 into two parts: one with *k* elements as the eigenvalues corresponding to common eigenvectors (denoted as 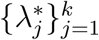) and the other one consisting of *p* − *k* elements (denoted as 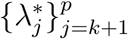). For *i* = 1, …, *n*, we independently sample *λ*_*ij*_ from a chi-squared distribution with degrees of freedom 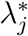 and construct 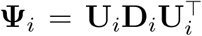, where **U**_*i*_ is an independent sample from the space {**U** : **U^T^U** = **I_*n*−*k*_, U^T^Γ** = **0**_(*n*−*k*)*×k*_} and **D**_*i*_ = diag{*λ*_*i*(*k*+1)_, …, *λ*_*ip*_}. Then we construct **Σ**_*i*_ = **Γ** diag{*λ*_*i*1_, …, *λ*_*ik*_}**Γ**^T^ + **Ψ**_*i*_ and {**y**_*it*_, *t* = 1, …, *T*} are independently sampled from *𝒩* (**0, Σ**_*i*_). If {*y*_*it*_} are not Gaussian distributed, we modify the above procedure by letting 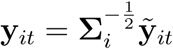, where 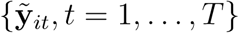 are independently sampled from a multivariate-Gamma distribution with mean **0**, variance **I**_*p*_ and skewness 10**I**_*p*_. If CPC-associated eigenvalues are the largest *k* eigenvalues across matrices, we set 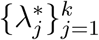 to be the largest *k* numbers in 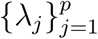.

### 4.2 Simulation results

Simulation results are summarized in Figure 2 and Tables 1 and 2 for simulations 1, 2 and 3 respectively.

**Table 1.** The accuracy of Algorithm 1 and the sequential testing procedure under different settings with p = 20.

| | Distribution | Average accuracy of $\hat{\Gamma}$ | | | Average $\hat{k}$ | | |
| --- | --- | --- | --- | --- | --- | --- | --- |
| | | $k = 1$ | $k = 10$ | $k = 20$ | $k = 1$ | $k = 10$ | $k = 20$ |
| $n = T = 15$ | Gaussian | 0.81 | 0.91 | 0.93 | - | - | - |
|  | Gamma | 0.67 | 0.71 | 0.83 | - | - | - |
| $n = T = 30$ | Gaussian | 0.95 | 0.97 | 0.95 | 1.20 | 9.68 | 18.92 |
|  | Gamma | 0.70 | 0.86 | 0.92 | 0.65 | 4.04 | 4.02 |
| $n = T = 100$ | Gaussian | 0.98 | 0.99 | 0.95 | 1.26 | 10.02 | 20.00 |
|  | Gamma | 0.96 | 0.98 | 0.95 | 1.01 | 9.50 | 13.73 |

**Table 2.** The accuracy of methods in estimating CPC under different scenarios. Semi-1: the proposed semiparametric method with k known. Semi-2: the proposed semiparametric method with k unknown. Flury: the Flury’s method. PVD: population value decomposition.

|  |  | Semi-1 | Semi-2 | Flury | PVD |
| --- | --- | --- | --- | --- | --- |
| Scenario 1 | $p = 20$ | 1.00 | 1.00 | 1.00 | 0.99 |
| | $p = 100$ | 1.00 | 1.00 | 1.00 | 0.99 |
| Scenario 2 | $p = 20$ | 0.99 | 0.96 | 0.26 | 0.50 |
| | $p = 100$ | 0.99 | 0.93 | 0.24 | 0.20 |
| Scenario 3 | $p = 20$ | 0.98 | 0.95 | 0.88 | 0.94 |
| | $p = 100$ | 1.00 | 0.95 | 0.91 | 0.95 |
| Scenario 4 | $p = 20$ | 0.99 | 0.99 | 0.99 | 0.95 |
| | $p = 100$ | 0.99 | 0.99 | 0.97 | 0.96 |

**Figure 2.**
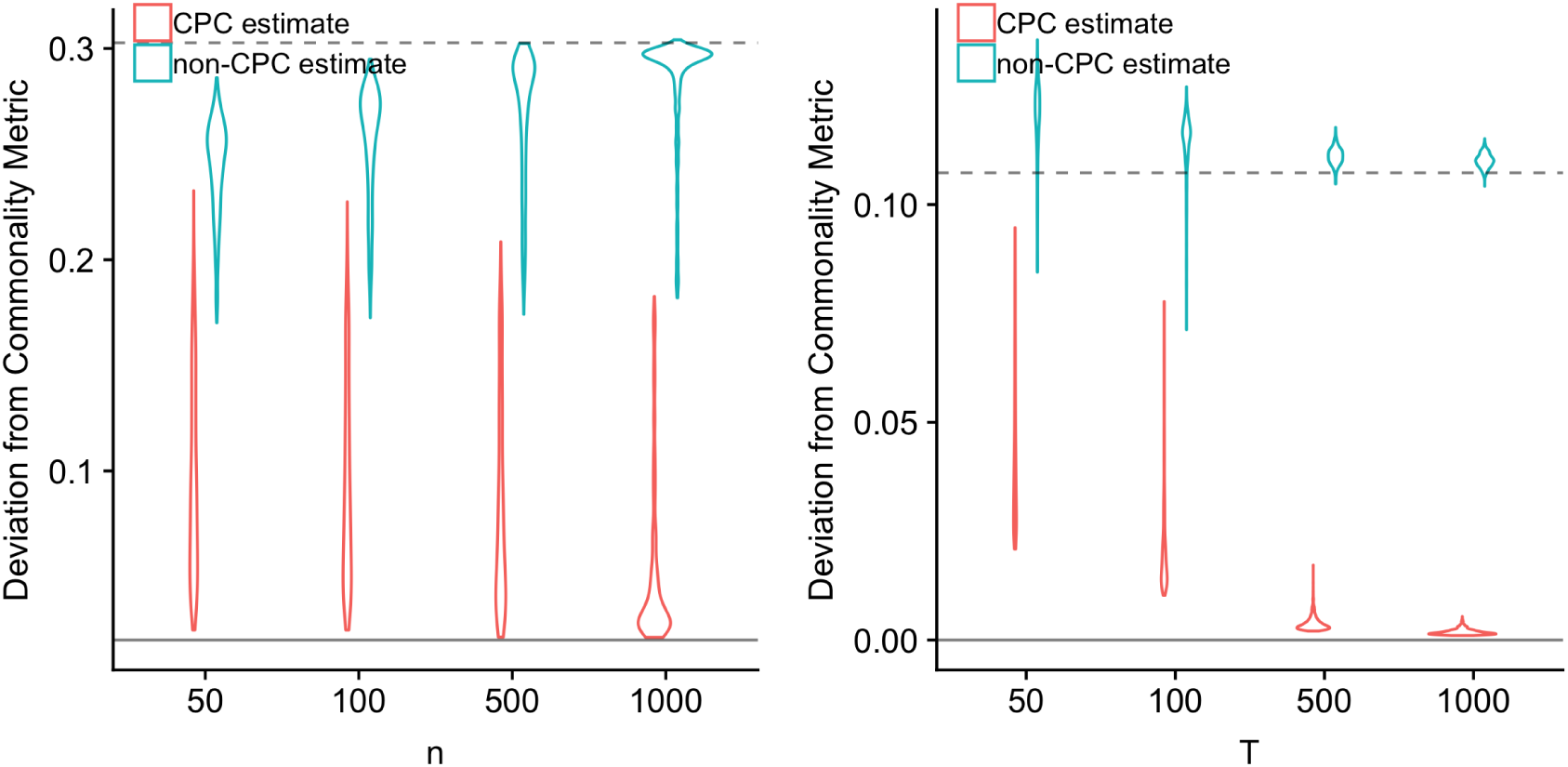
Distribution of the “Deviation from commonality” metric (3) as *n* (left panel) or *T* (right panel) goes to infinity for the last CPC estimate and the first non-CPC estimate. The solid line is the probability limit for the CPC estimate and the dashed line is the probability limit for the non-CPC estimate calculated from Theorem 2. The left panel demonstrates that the metric converges in probability to its limit, while the right panel shows that the metric converges almost surely.

Figure 2 shows that the deviation from commonality metric converges to its limit when *T* is fixed and *n* → ∞, and when *n* is fixed and *T* → ∞. This confirms the results of Theorem 2 and indicates that this metric can be used to distinguish CPC estimates and non-CPC estimates when either *n* or *T* is large.

Table 1 displays the performance of Algorithms 1 and 2 under the different simulation settings. When data are Gaussian distributed, both algorithms have high accuracy whenever *k, n, T* are small, medium or large. As *n* and *T* increases, the performance of both algorithms improves. When the sample size is small, Algorithm 1 still has a high accuracy in estimating **Γ**, even under the non-Gaussian distribution setting. In particular, when *n* = *T* = 15 *< p*, Algorithm 1 remains valid and has good accuracy, which demonstrates an advantage with small data. Since implementing Algorithm 2 requires *p* ≤ *T*, 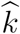 is not estimated when *n* = *T* = 15. As *k* increases, the accuracy of Algorithm 1 slightly increases, which confirms our discussion in Section 3.1. Under the Gamma data generating distribution, the algorithm to find *k* likely underestimates *k* when *k* is large. The reason for this is twofold. First, this algorithm is conservative for non-Gaussian data (as discussed in Section 3.2); second, when *k* is large, the number of null hypotheses to reject is large, which reduces the overall power. As a result, we recommend using Algorithm 2 for Gaussian distributed data. If *k* is large, one may not find all CPCs, but the identified ones are accurate.

Table 2 gives the comparison of our proposed method with *k* known or unknown to Flury’s method and PVD. In the first scenario, all methods perform well, as expected. In the other scenarios, our proposed method performs as good as or better than Flury’s method and PVD, even when the true number of CPCs is unknown. In Scenario 2, since both Flury’s method and PVD assume CPCs are associated with the largest eigenvalues for each matrix, their accuracy is much lower than our proposed method. In Scenario 3, all four methods have modest accuracy, but our proposed method with *k* unknown, Flury’s method and PVD have lower accuracy due to the non-Gaussian distribution. In contrast, our proposed method with *k* known remains highly accurate, since it is semiparametric. In Scenario 4, the size of data is limited compared to the dimension of matrices, resulting in small accuracy drops of all methods. However, our proposed method still has the least accuracy drop among all methods. In all scenarios, our proposed method outperforms Flury’s method and PVD, even when the true number of CPCs is unknown.

## 5. Task fMRI data example

We apply the proposed semiparametric PCPC method to the Human Connectome Project (HCP) motor-task fMRI data. The HCP project studies the brain connectome, both structural and functional, of healthy adults. The data set includes *n* = 136 healthy young adults from the most recent S1200 release. Adapted from the experimental design in Buckner et al. (2011) and Thomas Yeo et al. (2011), the task fMRI consists of ten task blocks including two tongue movement blocks, four hand movement blocks (two left and two right) and four foot movement blocks (two left and two right), as well as three 15-second fixation blocks. In each movement task block, a three-second visual cue was first presented followed by a 12-second movement. Participants were instructed to follow the visual cue to either move their tongue, or tap their left/right fingers, or squeeze their left/right toes to map the corresponding motor areas. The tasks were randomly intermixed. Once the ordering was fixed, the task onsets are nearly consistent across participants. The fMRI data were collected for *T* = 284 time points (with repetition time = 0.72 seconds) and 264 brain regions and preprocessed following the HCP minimal preprocessing pipeline (Glasser et al., 2013). Furthermore, we fit an ARMA(1,1) model (Lindquist et al., 2008) for each brain region to remove temporal correlation. We extracted blood-oxygen-level dependent (BOLD) signals from *p* = 35 functional brain regions in the sensorimotor network (Power et al., 2011) and averaged over voxels within the 5 mm radius. According to the Doornik-Hansen test for multivariate normality by Doornik and Hansen (2008), there is no sufficient evidence to reject the null hypothesis that data are normally distributed (*p*-value 0.12). Since Flury’s method and PVD are not able to identify the CPCs associated with small eigenvalues and require prespecified *k*, we present the result from the proposed semiparametric method only. Results of Flury’s method and PVD letting *k* = *p* are given in the Supplementary Material, which differ from the results of the semiparametric method.

Among 35 CPC candidates, Algorithm 2 identifies 30 as CPCs, which explain 80% of the total variance of the average covariance matrix. Figure 1 in the Supplementary Material summarises the results of sequential testings. To explore the relationship between the identified CPCs and the motor tasks, we plot the average time course of each CPC estimate (i.e., 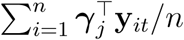 for *j* = 1, …,30) and compare it with task time bins. We also visualize brain regions with loading magnitude greater than 0.15 in a brain map. As a result, at least ten of the identified CPCs are related to tasks (no statistical test is performed) and a list of identified brain networks is provided in Table 3. Figure 3(A) presents an example of task-related CPC (CPC 18). In Figure 3(A), the average time course suggests a brain network of right hand movement and left foot movement, which is confirmed by the brain map. In this component, brain areas associated with motor control of the right hand yield high negative loadings (blue regions on the left hemisphere of the brain in Figure 3(A)); and regions associated with motor control of the left foot yield high positive loadings (red regions on the right hemisphere of the brain in Figure 3(A)). The lateral separation of the brain in terms of the loading sign suggests that during these motor tasks, the associated left and right hemispheres are functionally negatively correlated. Figure 3(B) presents an example of the CPC that is not related to the tasks. Even though the time course does not show a clear pattern, this CPC is concentrated in a region of the brain, which is modularized as a tongue region by Power et al. (2011). For the five components that are not identified as CPCs, two appear task-related and three do not. Since Algorithm 2 can be conservative when the true number of CPCs is large and the distribution is non-Gaussian based on simulation, some of these four CPC candidates may be CPCs. For all 35 CPC candidates, the average time course and the brain maps are provided in the Supplementary Material.

**Table 3.** Task-related brain networks identified by Algorithm 2.

| CPC No. | Variance explained | Brain network |
| --- | --- | --- |
| 3 | 2.0% | Hands, feet |
| 4 | 2.2% | Feet |
| 5 | 2.1% | Right hand |
| 10 | 2.3% | Right foot |
| 11 | 2.7% | Feet |
| 15 | 2.0% | Hands |
| 16 | 2.2% | Left hand |
| 18 | 2.1% | Right hand, left foot |
| 23 | 1.8% | Tongue, feet |
| 24 | 2.1% | Hands, feet |

**Figure 3.**
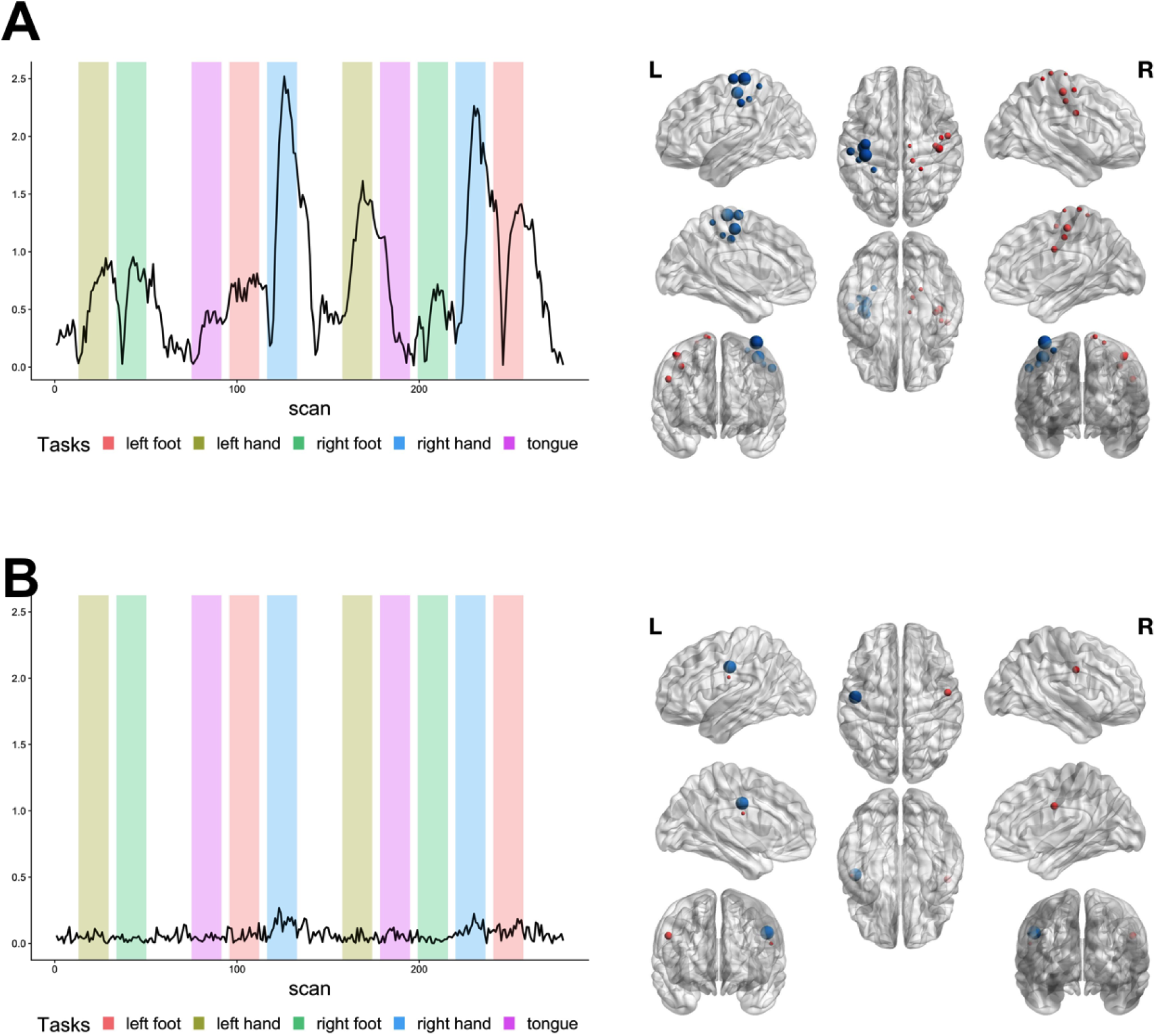
Average time course (left panel) and brain regions (right panel) of CPC 18 (upper panel) and CPC 9 (lower panel). In the left panel, each bin represents the time period of a task. In the right panel, each node is a brain region, with size standing for the absolute loading and color representing the sign of the loading (blue for negative and red for positive). Brain regions with absolute loading smaller than 0.1 are not shown in the figure.

Besides the analysis of the sensorimotor network, we ran Algorithm 2 on all brain regions (*p* = 264) and got 190 CPC estimates, which explain 50% of the total variance of the average covariance matrix. Among the 190 CPC estimates, 66 are associated with the default mode network, 15 are associated with the visual network, 12 are associated with the sensorimotor network and, 5 are associated with the frontoparietal network. Here we classify a CPC estimate as associated with a brain network if, among the regions with loadings greater than 0.1 in this CPC estimate, at least 25% come from the corresponding network. To compare the results from the sensorimotor-network analysis and whole-brain analysis, we extracted loadings corresponding to the sensorimotor network for each common eigenvector estimate in the whole-brain analysis. Among 30 CPC estimates of sensorimotor-network analysis, 12 are highly correlated (absolute value of inner product larger than 0.7) with some CPC candidates of whole-brain analysis, suggesting that the brain networks encoded by these CPCs retain when taking into account regions outside of the sensorimotor network.

## 6. Discussion

In this paper, we propose a semiparametric PCPC model and provide algorithms to identify CPCs with or without knowing the true number of CPCs. Furthermore, we prove the asymptotic consistency of our proposed estimators, even when the data generating distribution is non-Gaussian. In simulation studies, our estimator consistently outperforms Flury’s method and PVD and shows high accuracy if the number of CPCs is known. Applied to the motor-task fMRI data, our method identifies meaningful brain networks that match the current findings.

In PCPCA, a CPC may not be associated with the largest eigenvalues across all covariance matrices. For this reason, our proposed method allows for an arbitrary association between CPC and eigenvalues, which makes the model more flexible. One challenge resulting from this flexibility is to find *k*, the number of CPCs, since the signal of CPCs can be weak or inseparable from non-common principal components. Our proposed algorithm for finding *k* performs well under Gaussian distribution, but can be conservative if the underlying distribution is non-Gaussian or *k* is large. Furthermore, sequential hypothesis testing usually requires huge computational resources and can be slow for high-dimensional matrices. Hence, an efficient and robust method for finding *k* will be one future direction.

Our proposed method, as well as the literature, assumes *p*, the dimension of covariance matrices, is fixed. One exciting future direction could be finding solutions to handle data with large *p* but small *n* and *T*.

## Acknowledgement

Data were provided [in part] by the Human Connectome Project, WU-Minn Consortium (Principal Investigators: David Van Essen and Kamil Ugurbil; 1U54MH091657) funded by the 16 NIH Institutes and Centers that support the NIH Blueprint for Neuroscience Research; and by the McDonnell Center for Systems Neuroscience at Washington University. This work was supported by grants EB01503208, EB022911 and NS060910 from the National Institutes of Health.

## Supporting Information

Proofs of Theorems 1, 2, 3, and additional results of data application referenced in Section 5 are available with this paper at the Biometrics website on Wiley Online Library. The R code and data to reproduce the simulations and data application are available at https://github.com/BingkaiWang/Semi-parametric-PCPCA.

