## Supplementary Material for "Semiparametric Partial Common Principal Component Analysis for Covariance Matrices"

Summary of content in each section: we (A) prove our theoretical results; (B) provide additional results of the Semiparametric method on the fMRI data example; (C) present results of Flury's method and PVD on the fMRI data example.

### A Proofs

#### A.1 Proof of Theorem 1

To prove theorem 1, we first establish a rule of consistency of eigenvalues and eigenvectors.

**Lemma 1.** *For real positive definite symmetric matrices* $\mathbf{A}$ *and* $\mathbf{A}+\mathbf{E}_n, n=1,2,\dots,$ *let* $\alpha_j$ *and* $\alpha_{j,n}$ *denote the* $j$*-th largest eigenvalue of* $\mathbf{A}$ *and* $\mathbf{A}+\mathbf{E}_n$ *respectively for* $j=1,\dots,p$*. Then denote* $\mathbf{u}_j$ *and* $\mathbf{u}_{j,n}$ *as the corresponding unit eigenvector associated with* $\alpha_j$ *and* $\alpha_{j,n}$ *respectively. If* $||\mathbf{E}_n||_2 \xrightarrow{P} 0$*, where* $||\cdot||_2$ *is the matrix* $L_2$*-norm, and all eigenvalues of* $\mathbf{A}$ *are distinct, then* $\alpha_{j,n} \xrightarrow{P} \alpha_j$ *and* $|\mathbf{u}_j^\top \mathbf{u}_{j,n}| \xrightarrow{P} 1$ *for* $j=1,\dots,p$*.*

*Proof.* According to Weyl's inequality (Theorem 5.1 of Demmel, 1997), we have

$$|\alpha_j - \alpha_{j,n}| \leq ||\mathbf{E}_n||_2.$$

Since $||\mathbf{E}_n||_2 \xrightarrow{P} 0$, then $\alpha_{j,n} \xrightarrow{P} \alpha_j$. According to Theorem 5.4 of (Demmel, 1997), we have

$$\sin 2\theta_j \leq \frac{2||\mathbf{E}_n||_2}{\min_{i\neq j}|\alpha_i - \alpha_j|},$$

where $\theta_j$ is the acute angle between $\mathbf{u}_j$ and $\mathbf{u}_{j,n}$, i.e., $\theta_j = \arccos|\mathbf{u}_j^\top \mathbf{u}_{j,n}|$. Since $||\mathbf{E}_n||_2 \xrightarrow{P} 0$ and all eigenvalues of $\mathbf{A}$ are distinct, we have $\theta_j \xrightarrow{P} 0$ and hence $|\mathbf{u}_j^\top \mathbf{u}_{j,n}| \xrightarrow{P} 1$. □

*Proof of Theorem 1.* We first prove the asymptotics as $n \to \infty$. Define $\mathbf{\Lambda}^* = diag\{\lambda_1^*, \dots, \lambda_k^*\}$, $\mathbf{\Sigma}^* = (1 - \frac{1}{T})(\mathbf{\Gamma}\mathbf{\Lambda}^*\mathbf{\Gamma}^\top + \mathbf{\Psi}^*)$ and $\mathbf{E} = \frac{1}{n}\sum_{i=1}^n \mathbf{S}_i - \mathbf{\Sigma}^*$. We show $||\mathbf{E}||_2 \xrightarrow{P} 0$. For any $\varepsilon > 0$, since $||\mathbf{E}||_2 \le ||\mathbf{E}||_F \le \sqrt{p}||\mathbf{E}||_2$, we have

$$P(||\mathbf{E}||_2 > \varepsilon) \le P(||\mathbf{E}||_F > \sqrt{p}\varepsilon) \le \sum_{j,l=1}^{p} P(|E_{jl}| > \frac{\varepsilon}{\sqrt{p}}), \tag{1}$$

where $||\cdot||_F$ is the Frobenius norm and $E_{jl}$ is the an entry of $\mathbf{E}$ in its $j$-th row and $l$-th column. The inequality (1) shows that, to prove $||\mathbf{E}||_2 \xrightarrow{P} 0$, it suffices to show $E_{jl} \xrightarrow{P} 0$ for $j, k \in \{1, \dots, p\}$. To this end, we notice that Assumption A (2), (3) and (5) together imply that $\mathbf{y}_{it}$ has uniformly bounded fourth moment and $E[\mathbf{S}_i] = E[E[\mathbf{S}_i|\mathbf{\Sigma}_i]] = \mathbf{\Sigma}^*$. By week law of large numbers for triangular arrays (Durrett, 2019 Theorem 2.2.6), we get $\mathbf{E} \xrightarrow{P} 0$ element-wise. Finally, since $\boldsymbol{\gamma}_j, j = 1, \dots, k$ are eigenvectors of $\mathbf{\Sigma}^*$, Lemma 1 implies that a unit eigenvector $\boldsymbol{\gamma}_{l(j)}$ of $\frac{1}{n}\sum_{i=1}^n \mathbf{S}_i$ is consistent to $\boldsymbol{\gamma}_j$, where $l(j)$ is a function of $j$ independent of $n$ and $T$.

We then prove the asymptotics as $T \to \infty$. For fixed $n$, by Assumption B (4) and that $\mathbf{y}_{it}, t = 1, \dots, T$ are independent to each other, the strong law of large numbers implies that

$$\lim_{T\to\infty} \frac{1}{n}\sum_{i=1}^n \mathbf{S}_i \xrightarrow{a.s.} \frac{1}{n}\sum_{i=1}^n \mathbf{\Sigma}_i = \mathbf{\Gamma}\left(\frac{1}{n}\sum_{i=1}^n \mathbf{\Lambda}_i\right)\mathbf{\Gamma}^\top + \frac{1}{n}\sum_{i=1}^n \mathbf{\Psi}_i.$$

By Lemma 1 and assumption B (3), we get the desired result. □

#### A.2 Proof of Theorem 2

*Proof of Theorem 2.* The first part of this proof gives the asymptotics as $n \to \infty$ and the second part proves the asymptotics as $T \to \infty$.

**Part 1.** We introduce the following notations. Let $\hat{\boldsymbol{\gamma}}_j, j = 1, \dots, p$ be the $j$-th column of $\hat{\mathbf{\Gamma}}_{candi}$. Define $\mathbf{\Lambda}^* = diag\{\lambda_1^*, \dots, \lambda_k^*\}$ and $\mathbf{\Sigma}^* = (1 - \frac{1}{T})(\mathbf{\Gamma}\mathbf{\Lambda}^*\mathbf{\Gamma}^\top + \mathbf{\Psi}^*)$. By denoting the non-zero eigenvalues of $\mathbf{\Psi}^*$ as $\lambda_{k+1}^*, \dots, \lambda_p^*$ and the corresponding unit eigenvectors as $\boldsymbol{\gamma}_{k+1}, \dots, \boldsymbol{\gamma}_p$, we can write the eigenvalues of $\mathbf{\Sigma}^*$ as $(\lambda_1^*, \dots, \lambda_p^*)$ and the eigenbasis of $\mathbf{\Sigma}^*$ as $\mathbf{\Gamma}^* = (\boldsymbol{\gamma}_1, \dots, \boldsymbol{\gamma}_p)$. Without loss of generality, we can reorder columns of $\hat{\mathbf{\Gamma}}_{candi}$ such that $\hat{\boldsymbol{\gamma}}_j$ is consistent to $\boldsymbol{\gamma}_j$ for $j = 1, \dots, p$ according to Theorem 1. Furthermore, since $\hat{\boldsymbol{\gamma}}_j$ and $-\hat{\boldsymbol{\gamma}}_j$ represent the same eigenvector, without loss of generality, we can enforce the first non-zero element of $\hat{\boldsymbol{\gamma}}_j$ and $\boldsymbol{\gamma}_j$ to have the same sign, which implies that $\hat{\boldsymbol{\gamma}}_j - \boldsymbol{\gamma}_j \xrightarrow{P} 0$.

We first show that $\frac{1}{n}\sum_{i=1}^{n}[(\hat{\boldsymbol{\gamma}}_j^\top \mathbf{S}_i \hat{\boldsymbol{\gamma}}_l)^2 - (\boldsymbol{\gamma}_j^\top \mathbf{S}_i \boldsymbol{\gamma}_l)^2] \xrightarrow{P} 0$ for any $j, l \in \{1, \ldots, p\}$. It suffices to show $\frac{1}{n}\sum_{i=1}^{n}(\hat{\boldsymbol{\gamma}}_j - \boldsymbol{\gamma}_j)^\top \mathbf{S}_i \hat{\boldsymbol{\gamma}}_l \hat{\boldsymbol{\gamma}}_j^\top \mathbf{S}_i \hat{\boldsymbol{\gamma}}_l \xrightarrow{P} 0$. This is implied by the following derivation:

$$\begin{aligned}
\left|\frac{1}{n}\sum_{i=1}^{n}(\hat{\boldsymbol{\gamma}}_j - \boldsymbol{\gamma}_j)^\top \mathbf{S}_i \hat{\boldsymbol{\gamma}}_l \hat{\boldsymbol{\gamma}}_j^\top \mathbf{S}_i \hat{\boldsymbol{\gamma}}_l\right| &= \left|\frac{1}{n}\sum_{i=1}^{n} o_p(1) 1^\top \mathbf{S}_i \hat{\boldsymbol{\gamma}}_l \hat{\boldsymbol{\gamma}}_j^\top \mathbf{S}_i \hat{\boldsymbol{\gamma}}_l\right| \\
&\le o_p(1)\frac{1}{n}\sum_{i=1}^{n}\left|1^\top \mathbf{S}_i \hat{\boldsymbol{\gamma}}_l \hat{\boldsymbol{\gamma}}_j^\top \mathbf{S}_i \hat{\boldsymbol{\gamma}}_l\right| \\
&\le o_p(1)\frac{1}{n}\sum_{i=1}^{n}||\mathbf{S}_i||_2^2 \\
&= o_p(1)O_p(1) \\
&= o_p(1),
\end{aligned}$$

where $o_p(1)$ denotes a sequence of random vectors that converges to zero in probability and $O_p(1)$ represents a sequence that is bounded in probability. The second inequality above uses the fact that $\mathbf{y}_{it}$ has uniformly bounded fourth moment, which is implied by assumption A (2),(3) and (5).

We then show $\frac{1}{n}\sum_{i=1}^{n}(\boldsymbol{\gamma}_j^\top \mathbf{S}_i \boldsymbol{\gamma}_l)^2 \xrightarrow{P} \frac{1}{T}\lambda_j^* \lambda_l^*$ for any $j \in \{1, \ldots, k\}$ and $l \in \{1, \ldots, p\}$. Denoting

$$\boldsymbol{\Gamma}^{*t} \boldsymbol{\Sigma}_i \boldsymbol{\Gamma}^* = \begin{bmatrix} v_i^{(11)} & \ldots & v_i^{(p1)} \\ \vdots & \ddots & \vdots \\ v_i^{(p1)} & \ldots & v_i^{(pp)} \end{bmatrix},$$

then we have $v_i^{(jl)} = 0$ for $j \le k$ and $j \ne l$, and $v_i^{(jj)} = \lambda_{ij}$ for $j \le k$, given the PCPC model. By assumption A (5), for $j \le k$, we have

$$E[(\boldsymbol{\gamma}_j^\top \mathbf{S}_i \boldsymbol{\gamma}_l)^2] = E[E[(\boldsymbol{\gamma}_j^\top \mathbf{S}_i \boldsymbol{\gamma}_l)^2 | \boldsymbol{\Sigma}_i]] = \frac{1}{T}E[v_i^{(jj)} v_i^{(ll)} + 2 v_i^{(jl)2}] = \frac{1}{T}E[v_i^{(jj)} v_i^{(ll)}] = \frac{1}{T}E[\lambda_{ij} v_i^{(ll)}].$$

Since $v_i^{(ll)}$ does not involve $\lambda_{ij}$, assumption A (2) implies that $\lambda_{ij}$ is independnet of $v_i^{(ll)}$ and hence $E[(\boldsymbol{\gamma}_j^\top \mathbf{S}_i \boldsymbol{\gamma}_l)^2] = \frac{1}{T}E[\lambda_{ij}]E[v_i^{(ll)}] = \frac{1}{T}\lambda_j^* E[v_i^{(ll)}]$. Because the fourth moment of $\mathbf{y}_{it}$ is uniformly bounded, by weak law of triangular arrays (Durrett, 2019 Theorem 2.2.6) and continuous mapping theorem, $\frac{1}{n}\sum_{i=1}^{n}\{(\boldsymbol{\gamma}_j^\top \mathbf{S}_i \boldsymbol{\gamma}_l)^2 - \frac{1}{T}\lambda_j^* E[v_i^{(ll)}]\} \xrightarrow{P} 0$. If $l \le k$, then $v_i^{(ll)} = \lambda_{il}$ and hence $E[v_i^{(ll)}] = \lambda_l^*$. If $l > k$, Assumption A (2) and (3) imply that $\frac{1}{n}\sum_{i=1}^{n} E[v_i^{(ll)}] \xrightarrow{P} \lambda_l^*$. As a result, $\frac{1}{n}\sum_{i=1}^{n}(\boldsymbol{\gamma}_j^\top \mathbf{S}_i \boldsymbol{\gamma}_l)^2 \xrightarrow{P} \frac{1}{T}\lambda_j^* \lambda_l^*$ for $j \in \{1, \ldots, k\}$ and $l \in \{1, \ldots, p\}$.

On the other hand, if $j, l \in \{k+1, \ldots, p\}$, we show that $\frac{1}{n}\sum_{i=1}^{n}(\boldsymbol{\gamma}_j^\top \mathbf{S}_i \boldsymbol{\gamma}_l)^2 \xrightarrow{P} \frac{1}{T}[\lambda_j^* \lambda_l^* + 2Var(\boldsymbol{\gamma}_j^\top \boldsymbol{\Psi}_i \boldsymbol{\gamma}_l)]$. Similar to the previous case, we get $\frac{1}{n}\sum_{i=1}^{n}\{(\boldsymbol{\gamma}_j^\top \mathbf{S}_i \boldsymbol{\gamma}_l)^2 - \frac{1}{T}\lambda_j^*\lambda_l^* - \frac{2}{T}E[v_i^{(jl)2}]\} \xrightarrow{P} 0$. Since $j, l > k$ and $\boldsymbol{\gamma}_j$ and $\boldsymbol{\gamma}_l$ are different eigenvectors of $\boldsymbol{\Psi}^*$, then $v_i^{(jl)} = \boldsymbol{\gamma}_j^\top \boldsymbol{\Psi}_i \boldsymbol{\gamma}_l$ with $E[v_i^{(jl)}] = \boldsymbol{\gamma}_j^\top \boldsymbol{\Psi}^* \boldsymbol{\gamma}_l = 0$. Hence $E[v_i^{(jl)2}] = Var(\boldsymbol{\gamma}_j^\top \boldsymbol{\Psi}_i \boldsymbol{\gamma}_l)$. By assumption A (3), $Var(\boldsymbol{\gamma}_j^\top \boldsymbol{\Psi}_i \boldsymbol{\gamma}_l)$ is unchanged across $i$, which implies the desired result.

Finally, for $j \in \{1, \ldots, k\}$, by continuous mapping theorem and week law of large numbers,

$$\begin{aligned}
\text{Dev}(\{\mathbf{y}_{it}\}, \hat{\boldsymbol{\Gamma}}_{candi}, j) &= \frac{1}{n(p-1)} \sum_{l=1, l\neq j}^{p} \frac{\sum_{i=1}^{n}(\hat{\boldsymbol{\gamma}}_j^\top \mathbf{S}_i \hat{\boldsymbol{\gamma}}_l)^2}{\hat{\boldsymbol{\gamma}}_j^\top (\frac{1}{n}\sum_{i=1}^{n}\mathbf{S}_i)\hat{\boldsymbol{\gamma}}_j \hat{\boldsymbol{\gamma}}_l^\top (\frac{1}{n}\sum_{i=1}^{n}\mathbf{S}_i)\hat{\boldsymbol{\gamma}}_l} \\
&\xrightarrow{P} \frac{1}{(p-1)} \sum_{l=1, l\neq j}^{p} \frac{\frac{1}{T}\lambda_j^*\lambda_l^*}{\boldsymbol{\gamma}_j^\top \boldsymbol{\Sigma}^* \boldsymbol{\gamma}_j \boldsymbol{\gamma}_l^\top \boldsymbol{\Sigma}^* \boldsymbol{\gamma}_l} \\
&= \frac{1}{(p-1)} \sum_{l=1, l\neq j}^{p} \frac{\frac{1}{T}\lambda_j^*\lambda_l^*}{\lambda_j^*\lambda_l^*} \\
&= \frac{1}{T}.
\end{aligned}$$

For $j \in \{k+1, \ldots, p\}$, similarly,

$$\begin{aligned}
\text{Dev}(\{\mathbf{y}_{it}\}, \hat{\boldsymbol{\Gamma}}_{candi}, j) &= \frac{1}{n(p-1)} \sum_{l=1, l\neq j}^{p} \frac{\sum_{i=1}^{n}(\hat{\boldsymbol{\gamma}}_j^\top \mathbf{S}_i \hat{\boldsymbol{\gamma}}_l)^2}{\hat{\boldsymbol{\gamma}}_j^\top (\frac{1}{n}\sum_{i=1}^{n}\mathbf{S}_i)\hat{\boldsymbol{\gamma}}_j \hat{\boldsymbol{\gamma}}_l^\top (\frac{1}{n}\sum_{i=1}^{n}\mathbf{S}_i)\hat{\boldsymbol{\gamma}}_l} \\
&\xrightarrow{P} \frac{1}{(p-1)} \sum_{l=1, l\neq j}^{p} \frac{\frac{1}{T}\lambda_j^*\lambda_l^*}{\boldsymbol{\gamma}_j^\top \boldsymbol{\Sigma}^* \boldsymbol{\gamma}_j \boldsymbol{\gamma}_l^\top \boldsymbol{\Sigma}^* \boldsymbol{\gamma}_l} + \frac{1}{(p-1)} \sum_{l=k+1, l\neq j}^{p} \frac{2}{T} \frac{Var(\boldsymbol{\gamma}_j^\top \boldsymbol{\Psi}_i \boldsymbol{\gamma}_l)}{\boldsymbol{\gamma}_j^\top \boldsymbol{\Sigma}^* \boldsymbol{\gamma}_j \boldsymbol{\gamma}_l^\top \boldsymbol{\Sigma}^* \boldsymbol{\gamma}_l} \\
&= \frac{1}{(p-1)} \sum_{l=1, l\neq j}^{p} \frac{\frac{1}{T}\lambda_j^*\lambda_l^*}{\lambda_j^*\lambda_l^*} + \frac{1}{(p-1)} \sum_{l=k+1, l\neq j}^{p} \frac{2}{T} \frac{Var(\boldsymbol{\gamma}_j^\top \boldsymbol{\Psi}_i \boldsymbol{\gamma}_l)}{\lambda_j^*\lambda_l^*} \\
&= \frac{1}{T} + \frac{2}{T(p-1)} \sum_{l=k+1, l\neq j}^{p} \frac{Var(\boldsymbol{\gamma}_j^\top \boldsymbol{\Psi}_i \boldsymbol{\gamma}_l)}{\lambda_j^*\lambda_l^*}.
\end{aligned}$$

We define $C_j = \sum_{l=k+1, l\neq j}^{p} \frac{Var(\boldsymbol{\gamma}_j^\top \boldsymbol{\Psi}_i \boldsymbol{\gamma}_l)}{\lambda_j^*\lambda_l^*}$. It is straightforward that $C_j \geq 0$ and $C_j = 0$ if and only if $Var(\boldsymbol{\gamma}_j^\top \boldsymbol{\Psi}_i \boldsymbol{\gamma}_l) = 0$ for all $l > k$ and $l \neq j$. However, if $C_j = 0$, then $\boldsymbol{\gamma}_j^\top \boldsymbol{\Psi}_i \boldsymbol{\gamma}_l$ does not change across $i$ for all $l \neq j$, which means $\boldsymbol{\gamma}_j$ is a common eigenvector. This is contradicted with $j > k$. Hence $C_j > 0$. Then

$$\min_{l \in L_n} Err(\{\mathbf{y}_{it}\}, \hat{\boldsymbol{\Gamma}}_{candi}, l) \xrightarrow{P} \frac{1}{T} + \min_{l \in \{k+1, \ldots, p\}} C_l.$$

**Part 2.** Denote $\tilde{\mathbf{\Sigma}} = \frac{1}{n}\sum_{i=1}^{n}\mathbf{\Sigma}_i$ and let $(\mathbf{\Gamma}, \boldsymbol{\gamma}_{k+1}, \ldots, \boldsymbol{\gamma}_p)$ be the eigenbasis of $\tilde{\mathbf{\Sigma}}$. Similar to Part 1, without loss of generality, we assume that $\hat{\boldsymbol{\gamma}}_j \xrightarrow{a.s.} \boldsymbol{\gamma}_j$ for $j = 1, \ldots, p$.

By continuous mapping theorem and Theorem 1, as $T \to \infty$, for any $i = 1, \ldots, n$, $j, l \in \{1, \ldots, p\}$, we have

$$(\hat{\boldsymbol{\gamma}}_j^t \mathbf{S}_i \hat{\boldsymbol{\gamma}}_l)^2 \xrightarrow{a.s.} (\boldsymbol{\gamma}_j^t \mathbf{\Sigma}_i \boldsymbol{\gamma}_l)^2 \quad \text{and} \quad \hat{\boldsymbol{\gamma}}_j^t \left(\frac{1}{n}\sum_{i=1}^{n}\mathbf{S}_i\right)\hat{\boldsymbol{\gamma}}_j \xrightarrow{a.s.} \boldsymbol{\gamma}_j^t \left(\frac{1}{n}\sum_{i=1}^{n}\mathbf{\Sigma}_i\right)\boldsymbol{\gamma}_j.$$

If $\min\{j, l\} \le k$, according to the PCPC model, $\boldsymbol{\gamma}_j^t \mathbf{\Sigma}_i \boldsymbol{\gamma}_l = 0$ and hence $\text{Dev}(\{\mathbf{y}_{it}\}, \hat{\mathbf{\Gamma}}_{candi}, j) \xrightarrow{a.s.} 0$. If $\min\{j, l\} > k$, then

$$\text{Dev}(\{\mathbf{y}_{it}\}, \hat{\mathbf{\Gamma}}_{candi}, j) \xrightarrow{a.s.} \frac{1}{n(p-1)} \sum_{l=k+1, l\neq j}^{p} \frac{\sum_{i=1}^{n}(\boldsymbol{\gamma}_j^t \mathbf{\Sigma}_i \boldsymbol{\gamma}_l)^2}{\boldsymbol{\gamma}_j^t \tilde{\mathbf{\Sigma}} \boldsymbol{\gamma}_j \boldsymbol{\gamma}_l^t \tilde{\mathbf{\Sigma}} \boldsymbol{\gamma}_l} \ge 0.$$

The limit is 0 if and only if $\boldsymbol{\gamma}_j^t \mathbf{\Sigma}_i \boldsymbol{\gamma}_l = 0$ for all $i = 1, \ldots, n$. This, however, will imply that $\boldsymbol{\gamma}_j$ is a common eigenvector, which is contradicted with $j > k$. Hence the limit is greater than 0. Then

$$\min_{l \in L_n} Err(\{\mathbf{y}_{it}\}, \hat{\mathbf{\Gamma}}_{candi}, l) \xrightarrow{P} \min_{l \in \{k+1, \ldots, p\}} \frac{1}{n(p-1)} \sum_{j=k+1, j\neq l}^{p} \frac{\sum_{i=1}^{n}(\boldsymbol{\gamma}_j^t \mathbf{\Sigma}_i \boldsymbol{\gamma}_l)^2}{\boldsymbol{\gamma}_j^t \tilde{\mathbf{\Sigma}} \boldsymbol{\gamma}_j \boldsymbol{\gamma}_l^t \tilde{\mathbf{\Sigma}} \boldsymbol{\gamma}_l}.$$

□

#### A.3 Proof of Theorem 3

*Proof of Theorem 3.* Given the assumptions, the data generating distribution of $\{\mathbf{y}_{it}^{(\text{sim})}\}$ described in Section 3.2 is the same as the data generating distribution of $\{\mathbf{y}_{it}\}$. As a result, $\text{Dev}\,(\{\mathbf{y}_{it}^{(\text{sim})}\}, \hat{\mathbf{\Gamma}}_{\text{candi}}^{(\text{sim})}, j+1)$ calculated from each simulated data follow the distribution of $\text{Dev}\,(\{\mathbf{y}_{it}\}, \hat{\mathbf{\Gamma}}_{\text{candi}}, j+1)$. Since each simulated data are independent, as $m \to \infty$, the empirical distribution $\hat{F}_{j+1}$ converges to the distribution of $\text{Dev}\,(\{\mathbf{y}_{it}\}, \hat{\mathbf{\Gamma}}_{\text{candi}}, j+1)$ by the Glivenko-Cantelli theorem (Theorem 2.4.7 of Durrett, 2019). □

### B Additional results of the Semiparametric method on the fMRI data example

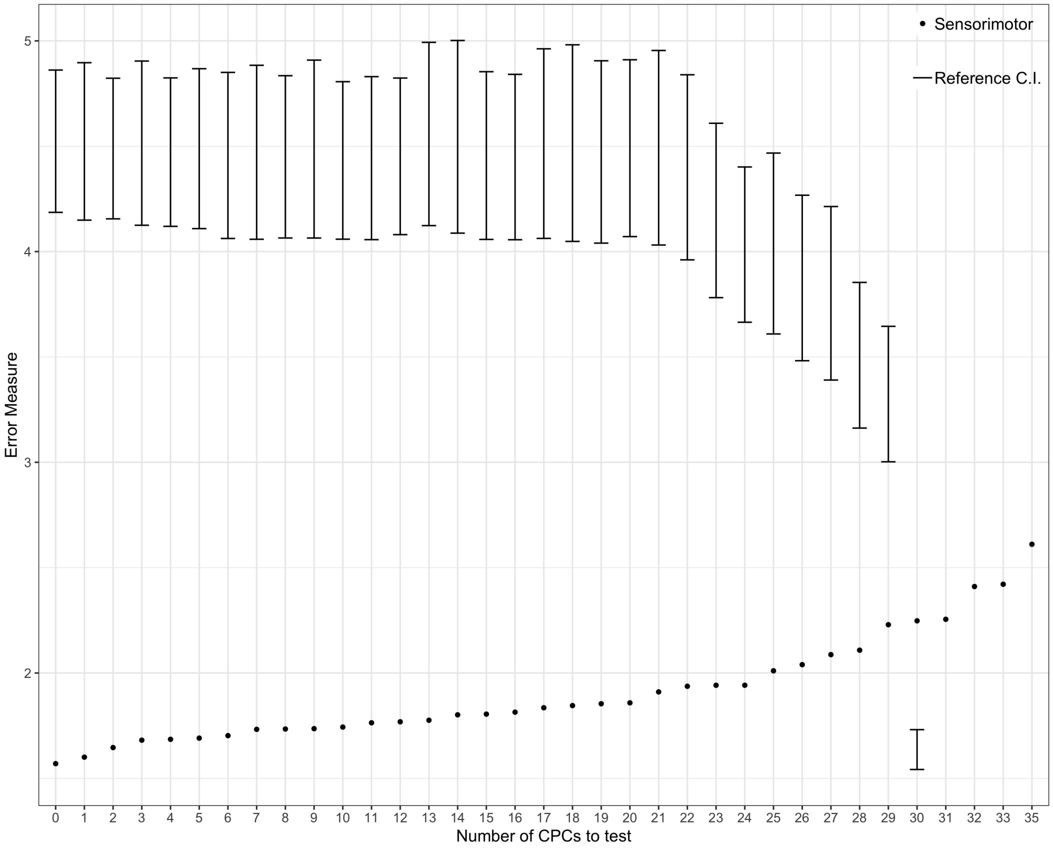


Figure 1: Summary of the sequential testing procedure. Each dot represents a CPC candidate calculated from the data set and each error bar represents the 95% confidence interval (CI) coverage derived from the simulation. When the dot is below the lower bound of the 95% CI, the null hypothesis is rejected and the next hypothesis is tested; otherwise the procedure stops.

#### B.1 Visualization of all CPCs

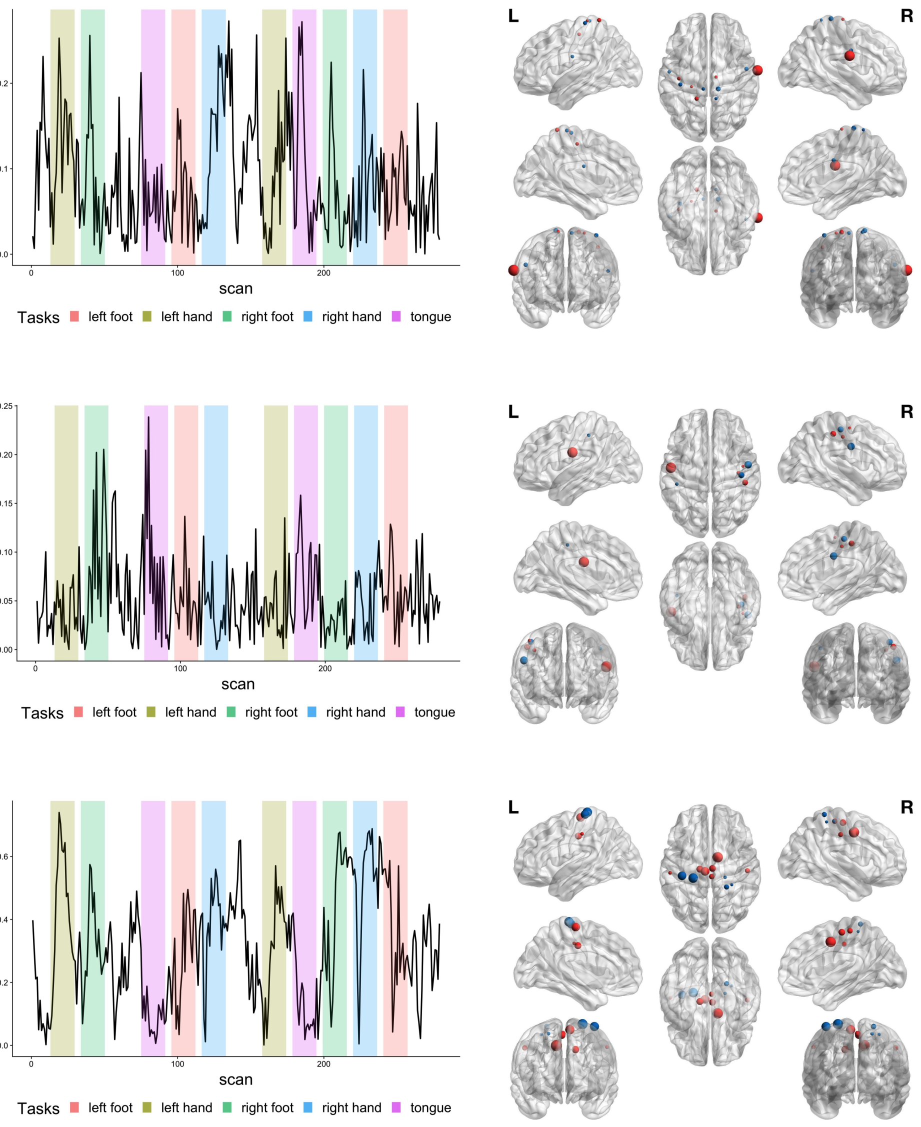


Figure 2: Average time course (left panel) and brain regions (right panel) of CPC 1 (upper panel), CPC 2 (middle panel) and CPC 4 (lower panel).

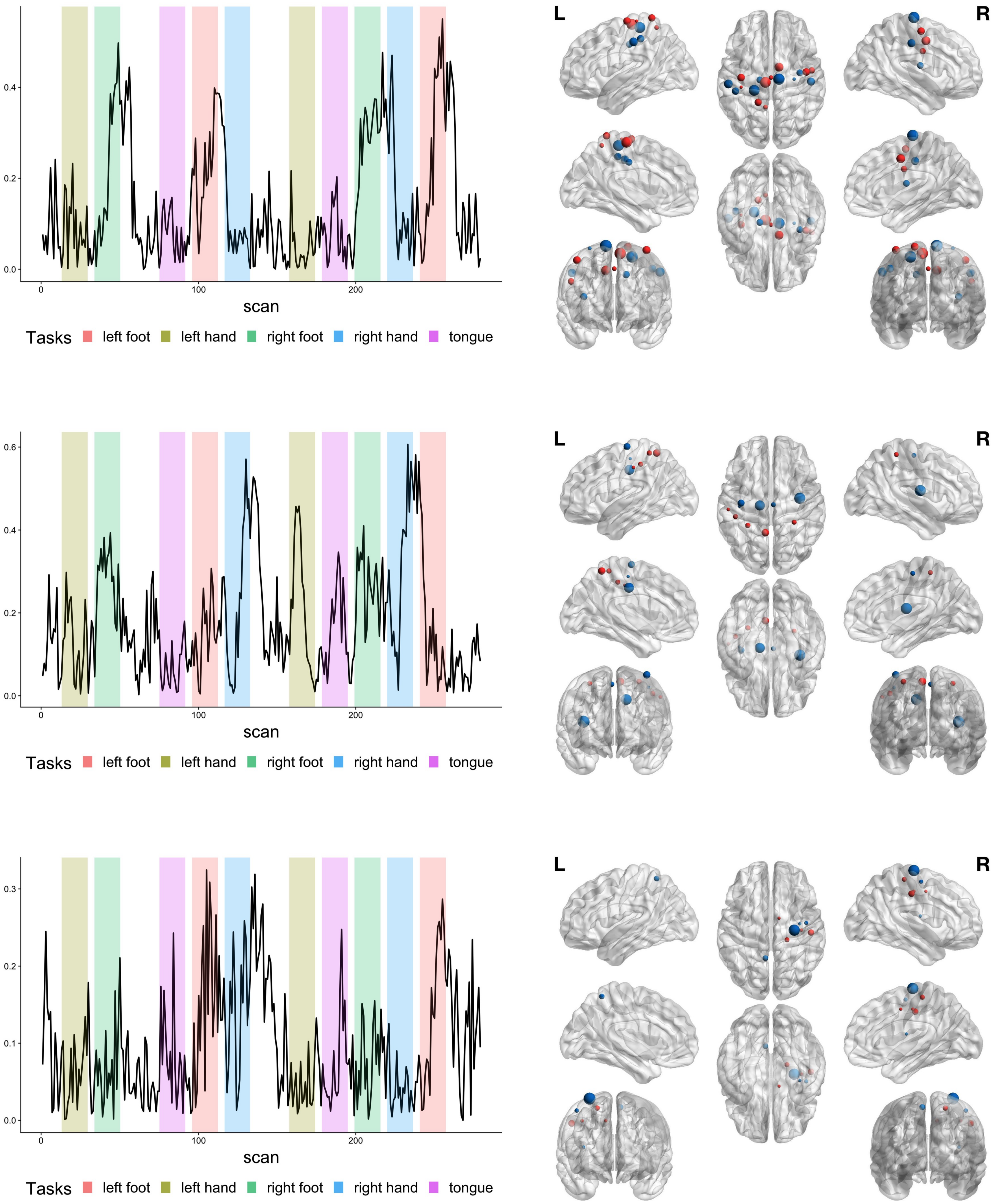


Figure 3: Average time course (left panel) and brain regions (right panel) of CPC 4 (upper panel), CPC 5 (middle panel) and CPC 6 (lower panel).

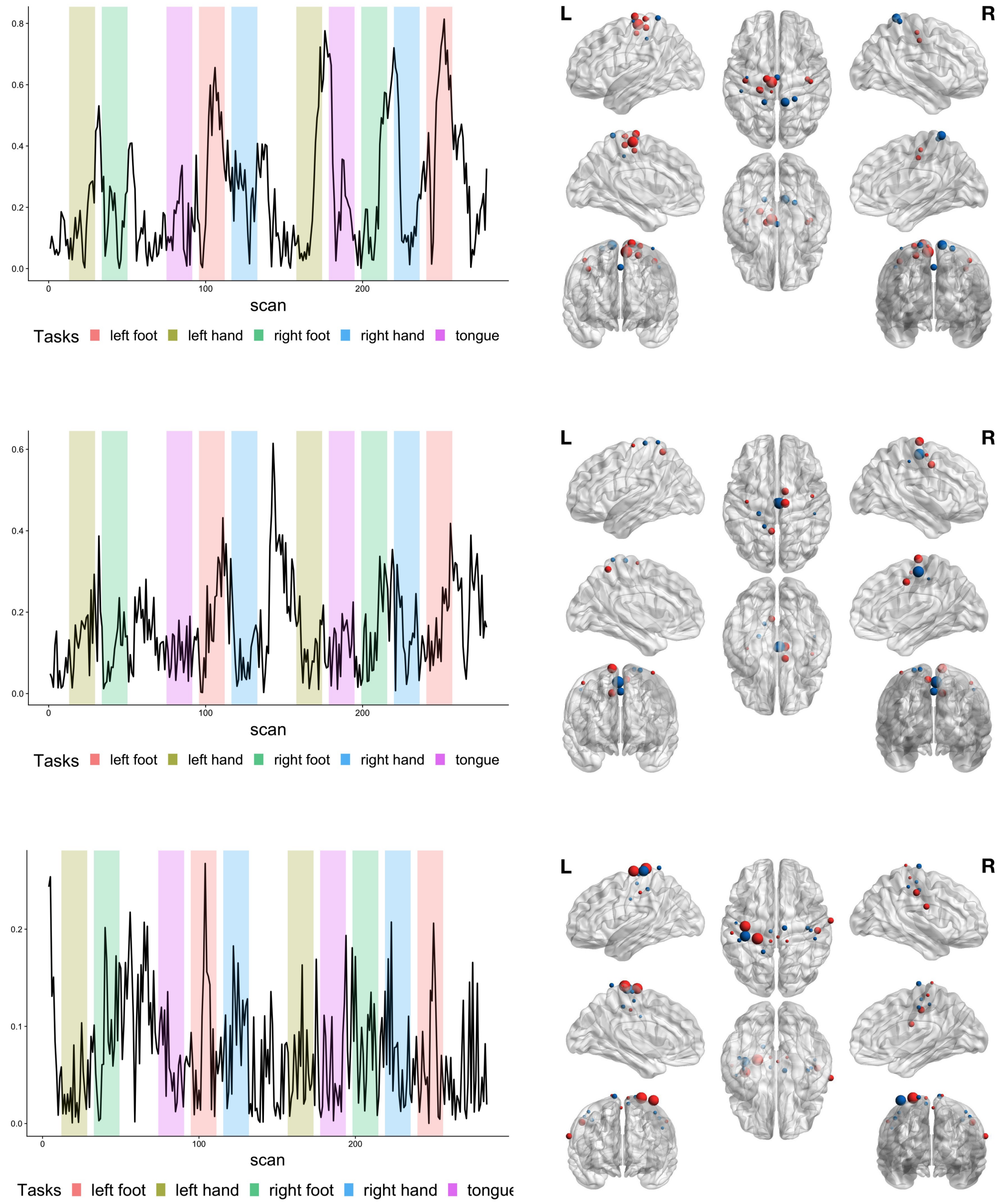


Figure 4: Average time course (left panel) and brain regions (right panel) of CPC 7 (upper panel), CPC 8 (middle panel) and CPC 9 (lower panel).

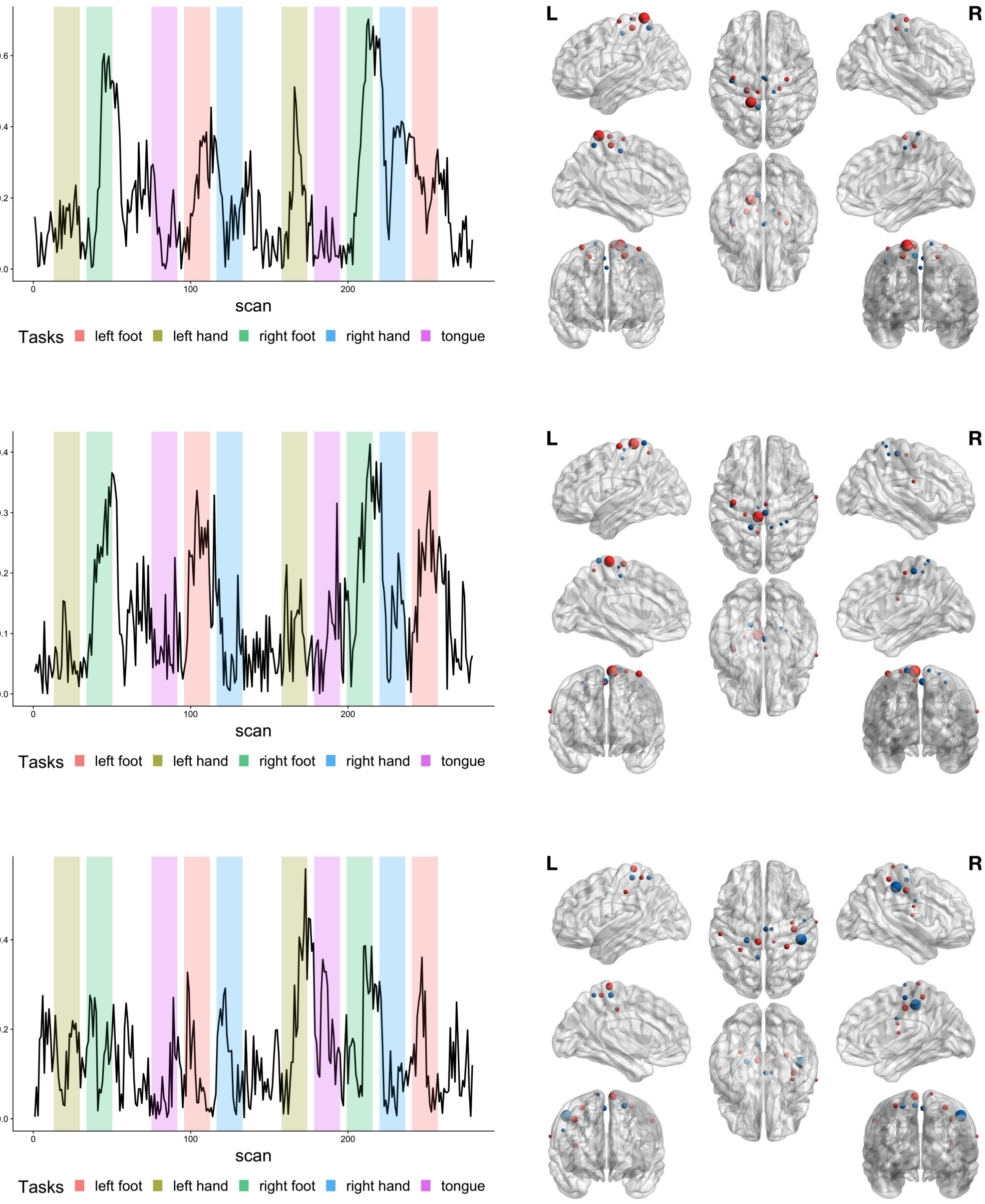


Figure 5: Average time course (left panel) and brain regions (right panel) of CPC 10 (upper panel), CPC 11 (middle panel) and CPC 12 (lower panel).

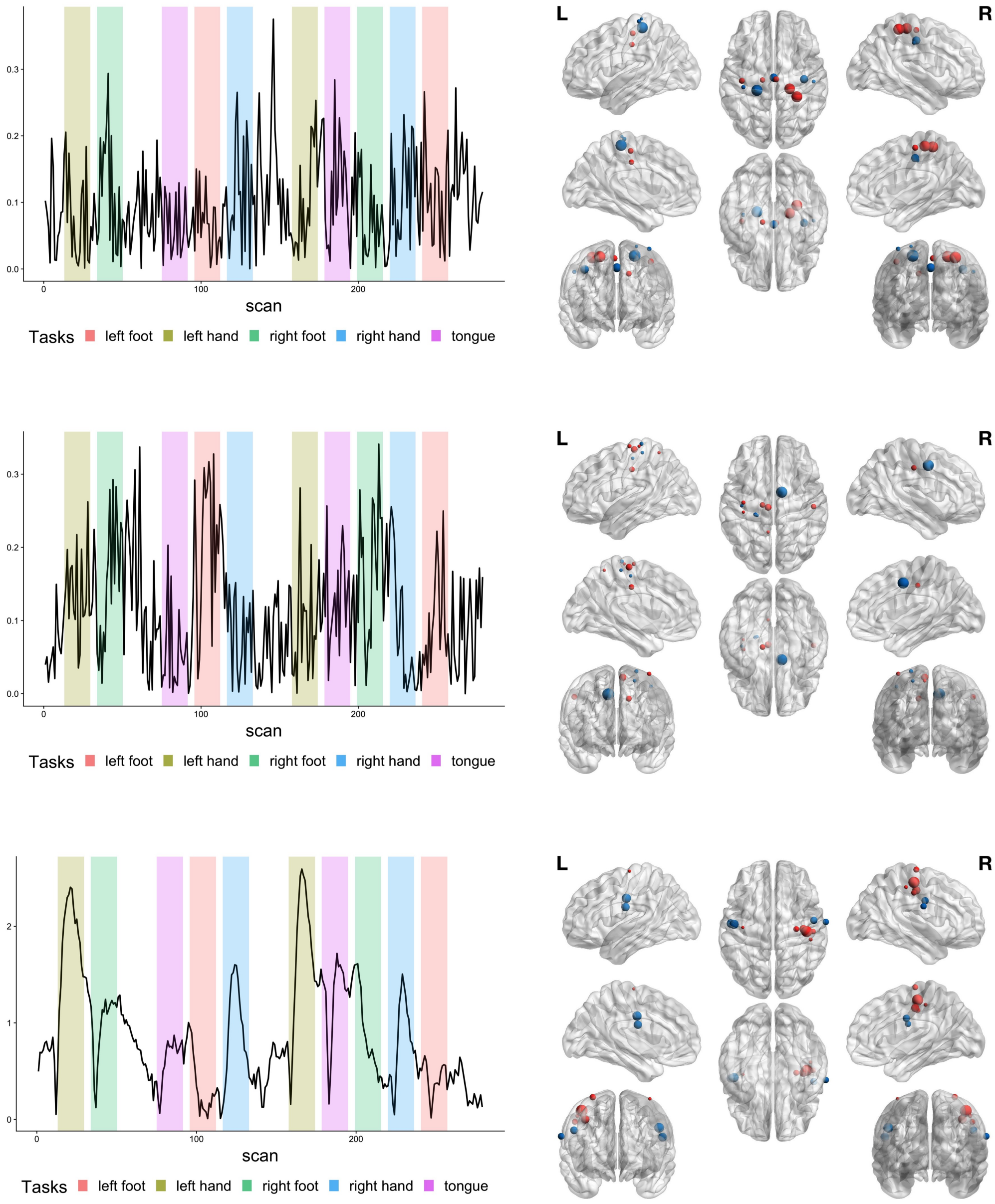


Figure 6: Average time course (left panel) and brain regions (right panel) of CPC 13 (upper panel), CPC 14 (middle panel) and CPC 15 (lower panel).

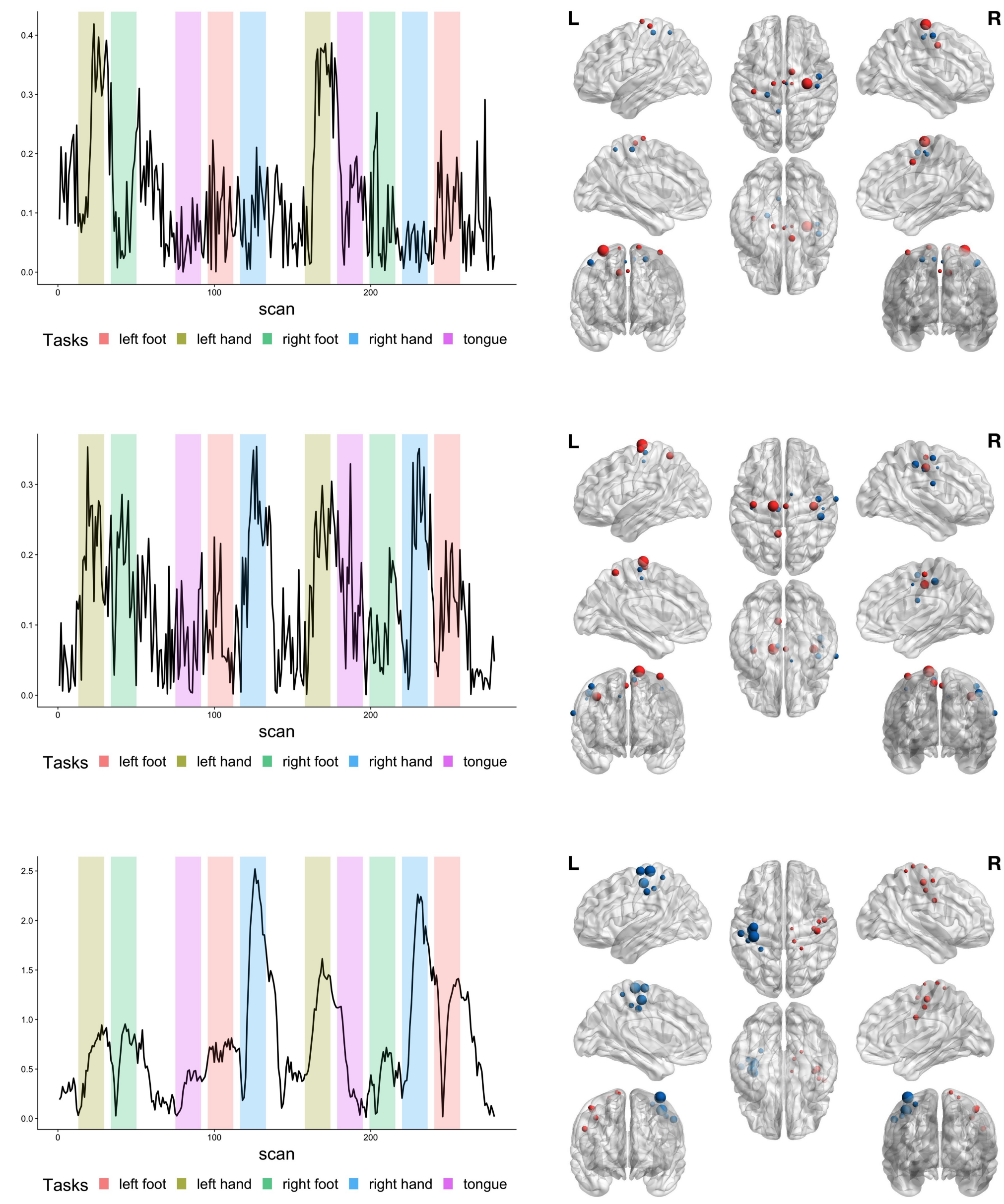


Figure 7: Average time course (left panel) and brain regions (right panel) of CPC 16 (upper panel), CPC 17 (middle panel) and CPC 18 (lower panel).

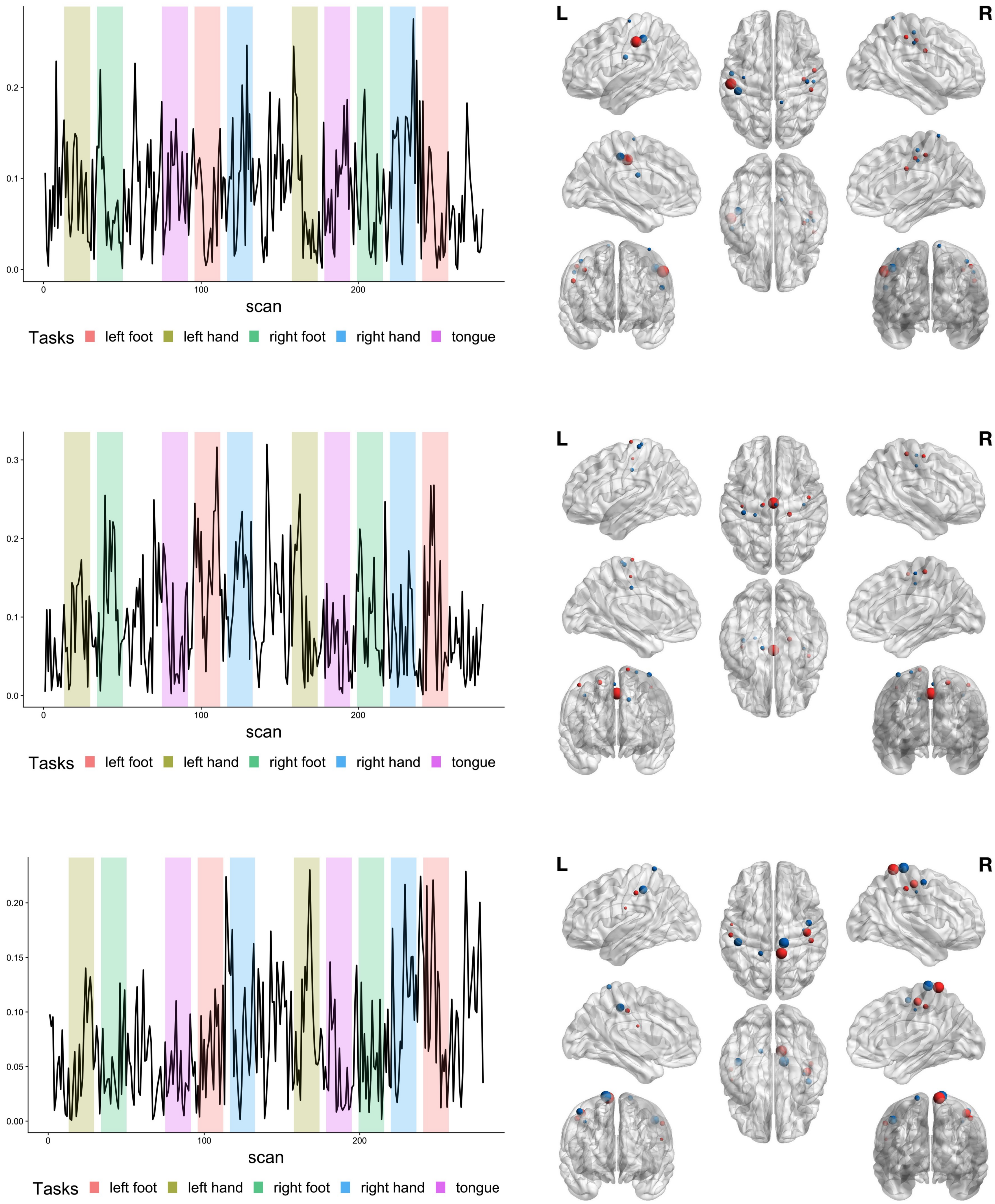


Figure 8: Average time course (left panel) and brain regions (right panel) of CPC 19 (upper panel), CPC 20 (middle panel) and CPC 21 (lower panel).

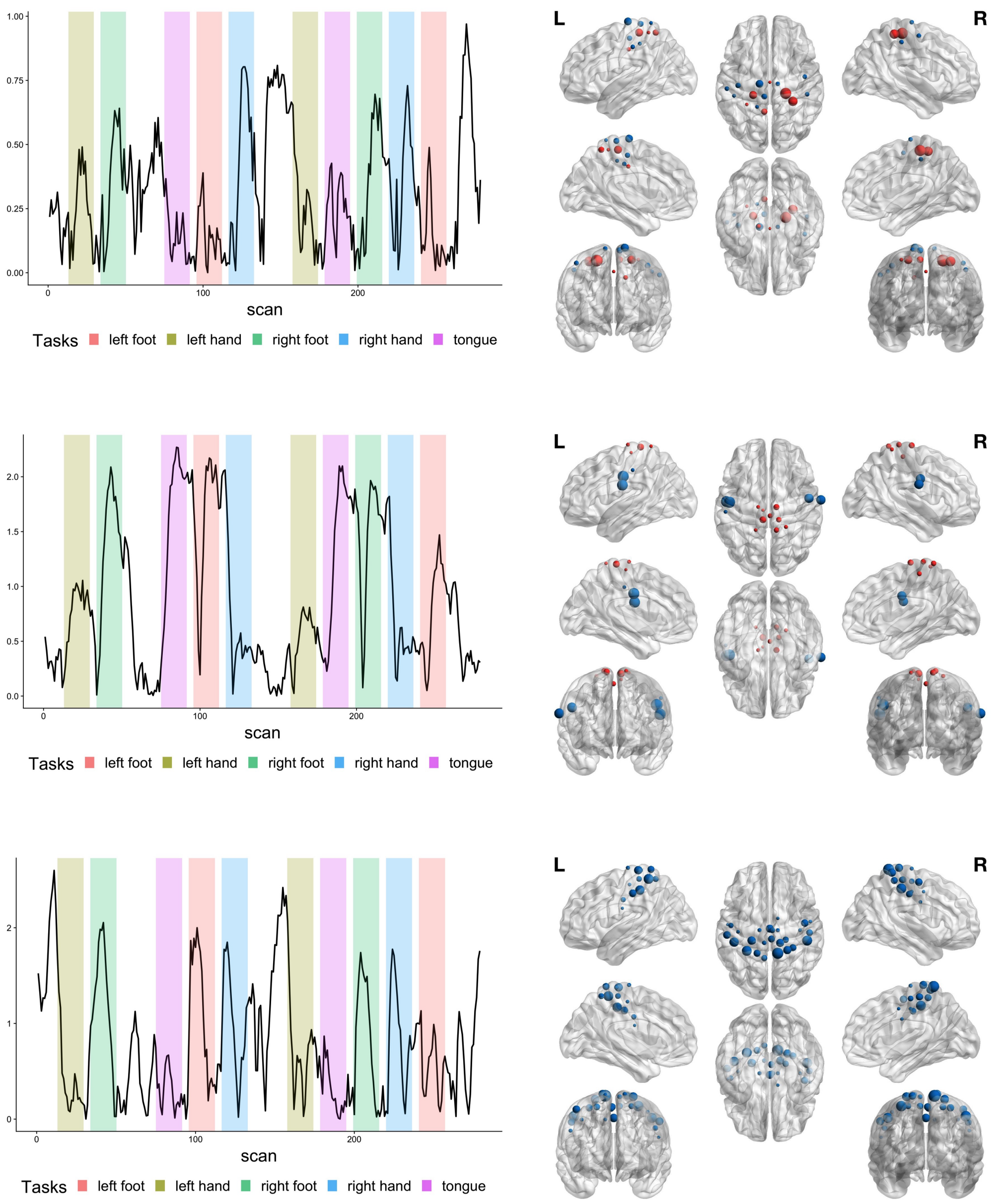


Figure 9: Average time course (left panel) and brain regions (right panel) of CPC 22 (upper panel), CPC 23 (middle panel) and CPC 24 (lower panel).

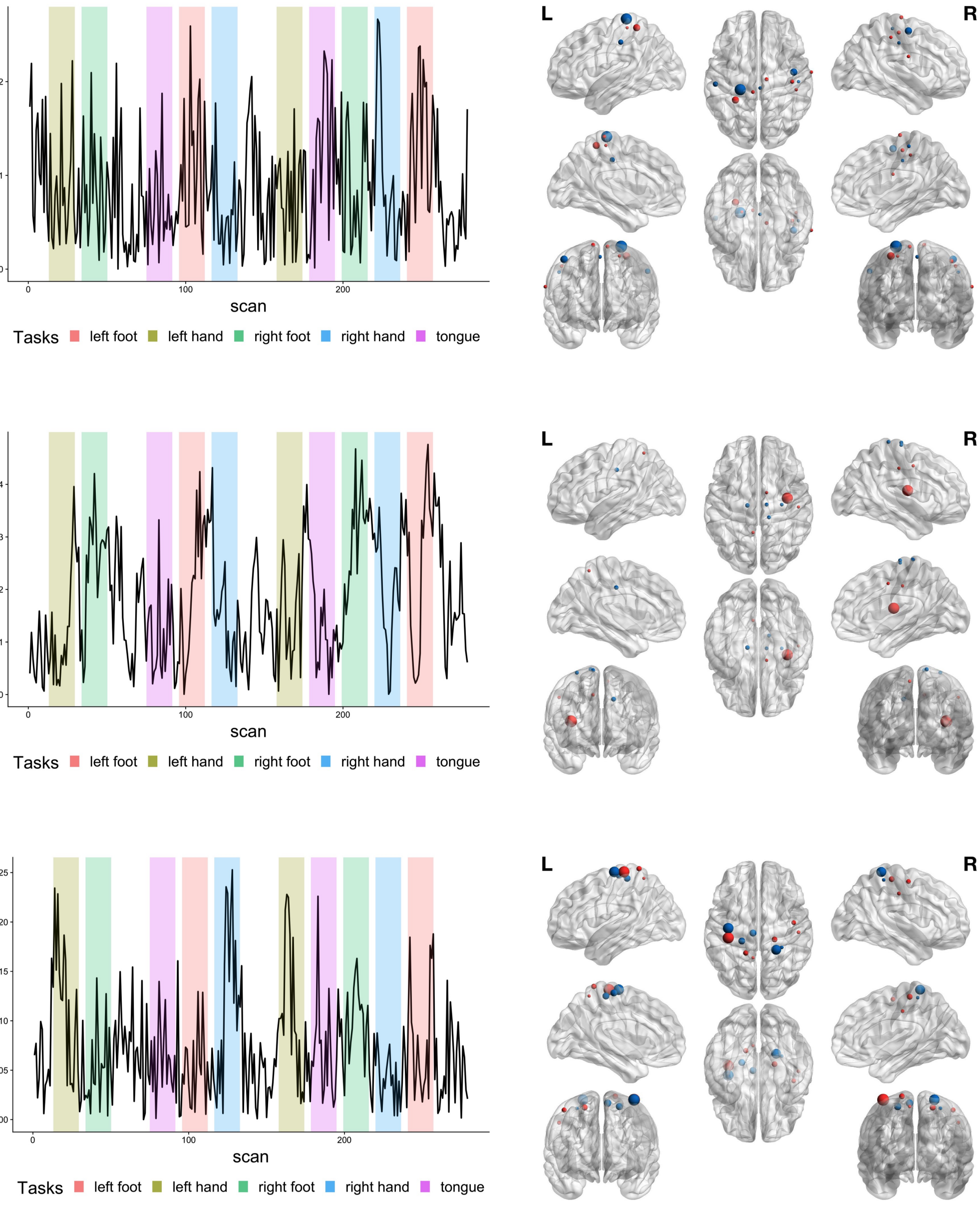


Figure 10: Average time course (left panel) and brain regions (right panel) of CPC 25 (upper panel), CPC 26 (middle panel) and CPC 27 (lower panel).

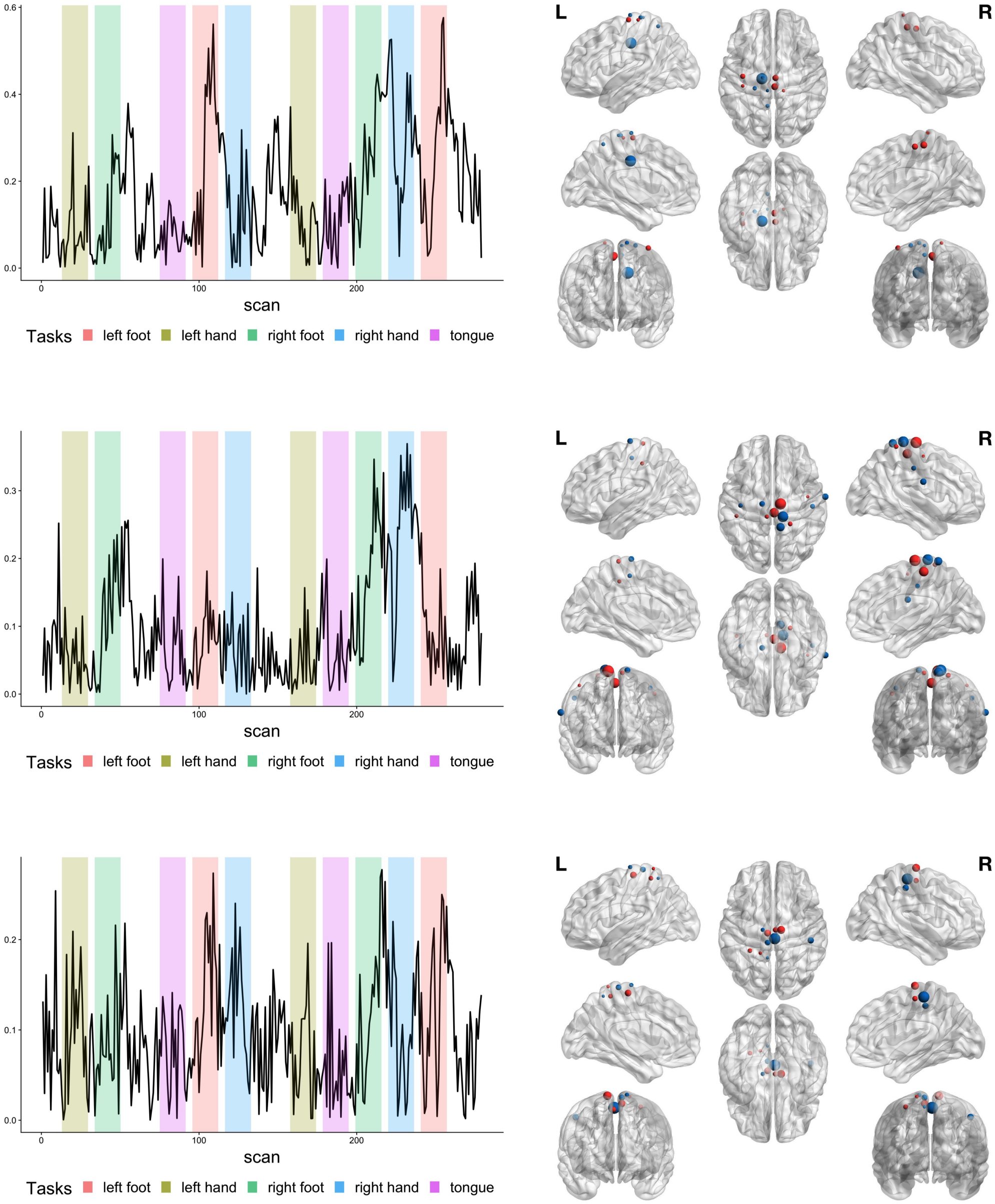


Figure 11: Average time course (left panel) and brain regions (right panel) of CPC 28 (upper panel), CPC 29 (middle panel) and CPC 30 (lower panel).

#### B.2 Visualization of non-CPCs

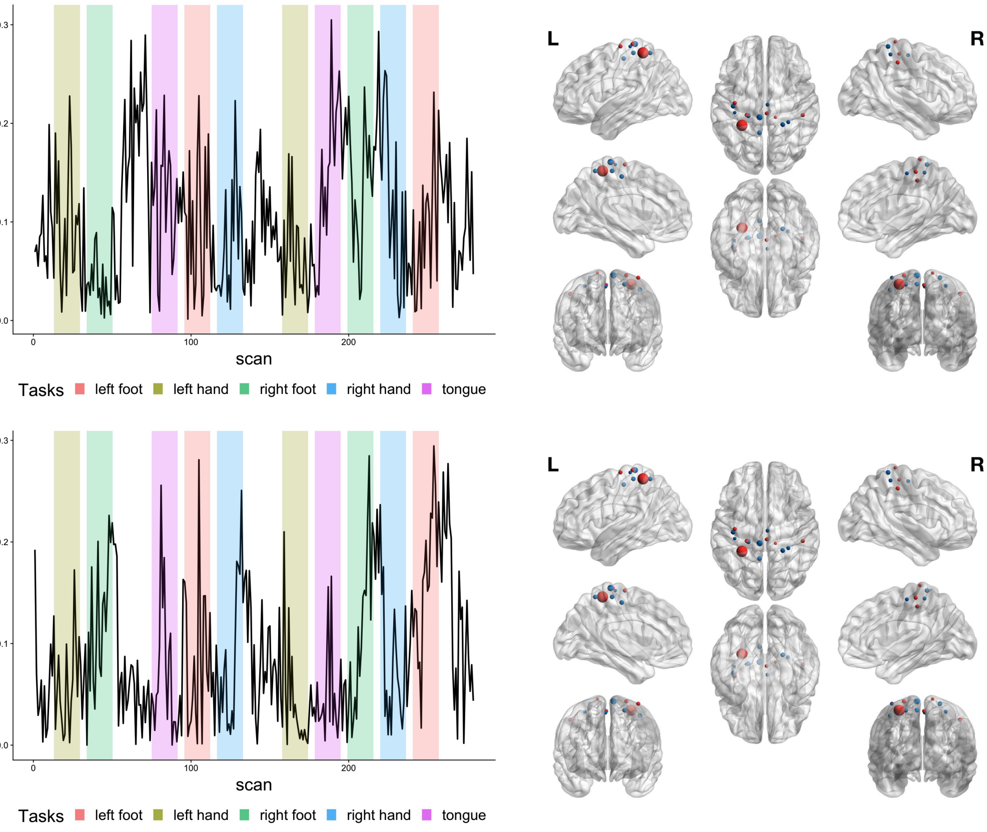


Figure 12: Average time course (left panel) and brain regions (right panel) of non-CPC 1 (upper panel) and non-CPC 2 (middle panel).

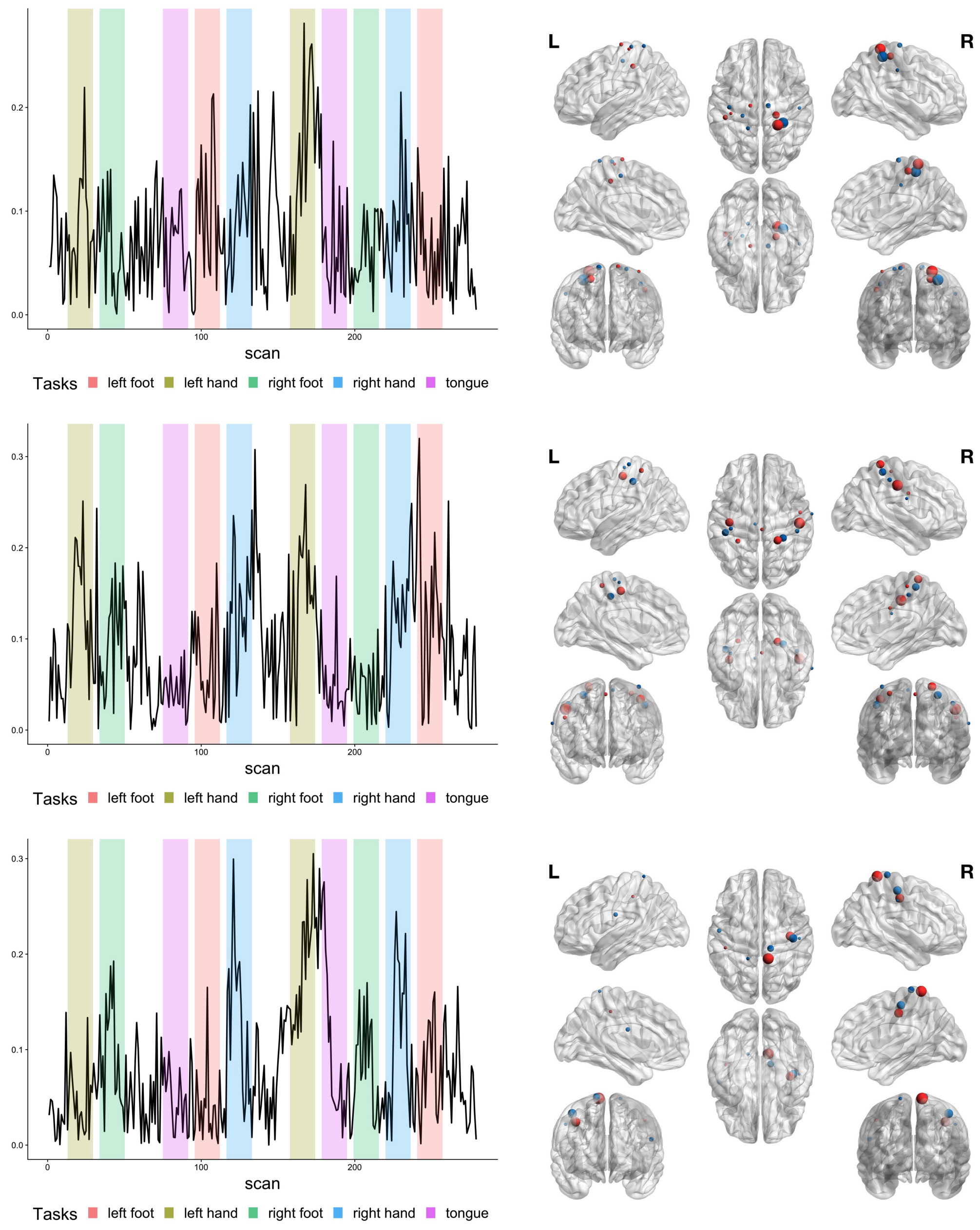


Figure 13: Average time course (left panel) and brain regions (right panel) of non-CPC 3 (upper panel), non-CPC 4 (middle panel) and non-CPC 5 (lower panel).

### C Results of Flury's method and PVD on the fMRI data example

Figures 14 and 15 summarize the CPCs identified by Flury's method and PVD respectively. Since neither method is able to find the number of CPC, we set $k = p$, which may resulting in false discovery of CPCs. Unlike our proposed method, CPCs are ranked according to their associated eigenvalues, and hence top CPCs may not be true CPCs. The time course of each CPC identified by Flury's method and PVD can be found at https://github.com/BingkaiWang/Semi-parametric-PCPCA.

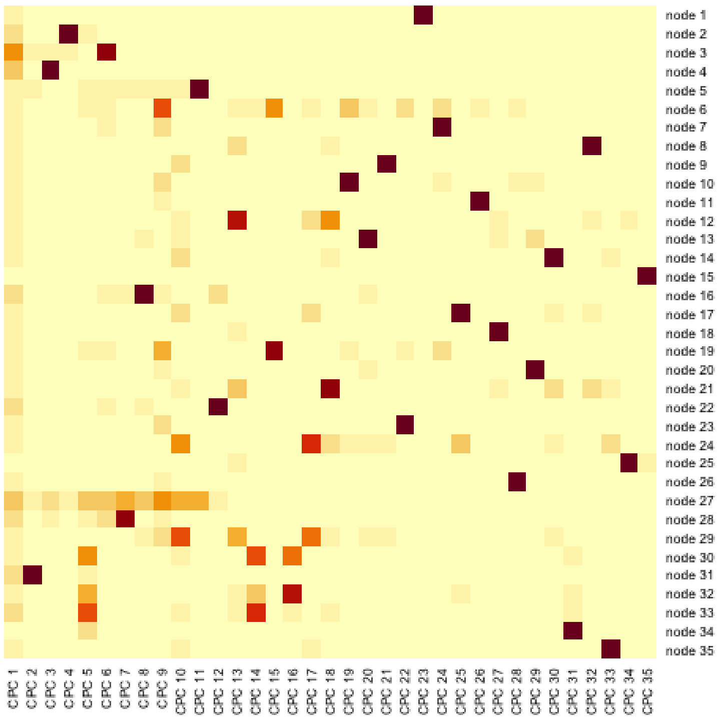


Figure 14: Summary of CPCs identified by the Flury's method. Each cell represents the absolute loading of a node of a CPC. Large (small) values are represented by white (red) color.

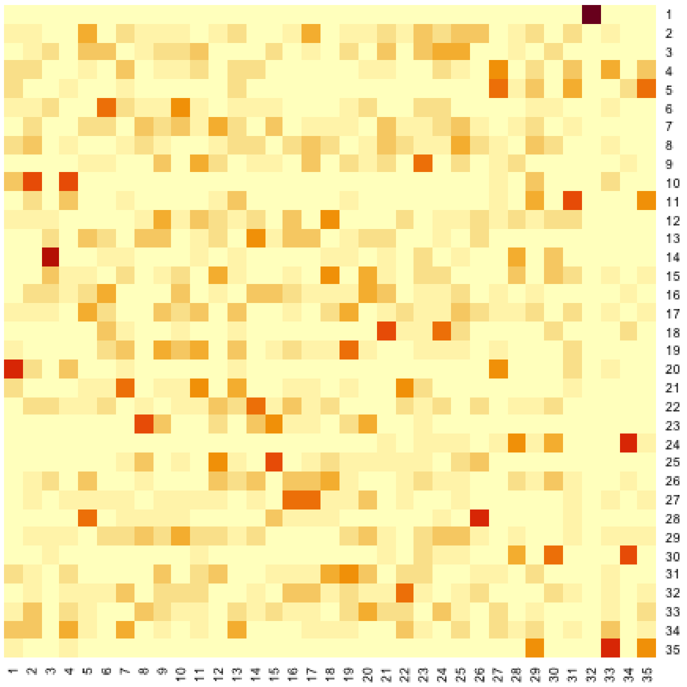

Figure 15: Summary of CPCs identified by PVD. Each cell represents the absolute loading of a node of a CPC. Large (small) values are represented by white (red) color.
